# A Mathematical Model of Reliability-Dependent Sensory Reweighting in Quiet Standing

**DOI:** 10.64898/2026.06.23.733972

**Authors:** Jun Kobayashi

## Abstract

Human quiet standing relies on vestibular, proprioceptive, and visual information whose contributions change with sensory context. Posturography characterizes this reweighting through responses to visual and support-surface perturbations, but a compact account of how sensory reliability propagates from state estimation to postural action remains incomplete. We develop a minimal quiet-standing model in which sensory reliability is encoded by channel-specific precision parameters that weight sensory prediction errors. A one-link inverted pendulum with intrinsic ankle stiffness estimates posture by minimizing an active-inference free-energy objective and selects ankle torque by minimizing the same objective under a restorative sensory goal. A fixed-point analysis predicts in closed form the perturbation-induced belief bias, each channel’s state-update contribution, the steady posture, and a linear stability condition from the precisions, goal dynamics, and intrinsic stiffness. Closed-loop simulations in 115 trials matched the predicted equilibria to within 0.001° wherever the loop settled, and the two conditions predicted unstable diverged or drifted. Reducing an unreliable channel’s precision eightfold reduced perturbation-driven posture shifts by 82%, with a matching decrease in that channel’s stateupdate contribution; intrinsic stiffness left this reduction essentially unchanged for visual and sensory-only perturbations but bounded it for physical support tilts; a global precision reduction did not reduce the shift; and a within-trial precision switch showed belief updating, not a separate action gain, carrying the change to ankle torque. The results give a compact mathematical account of postural sensory reweighting as relative, context-selective precision control in a closed multisensory loop.

## 1 Introduction

Adaptive postural control requires the nervous system to combine sensory information whose reliability changes with context. In quiet standing, visual, vestibular, and proprioceptive sensory signals do not contribute with fixed weights: standing on a firm surface with stable vision differs from standing with unreliable vision or a moving support surface. Human posturography has described this behavior as sensory reweighting, with perturbation-response curves changing as sensory reliability changes (Peterka, 2002; Peterka et al., 2018), and has characterized its time course after cues are added or removed (Assländer and Peterka, 2014, 2016). The computational question is how to express such reweighting as a control mechanism rather than a condition-specific gain schedule.

A useful mathematical account of this phenomenon should satisfy three requirements. It should (1) represent multiple sensory channels, (2) allow their reliability to vary with context, and (3) show how reliability changes affect both state estimation and postural action. Active inference provides one compact way to implement such an account. In continuous active inference, perception updates beliefs by reducing sensory and dynamical prediction errors, and action changes the world so that sensory input conforms to preferred predictions (Friston, 2010; Friston et al., 2010; Adams et al., 2013). Because perception and action minimize the same free-energy functional, active inference provides a closed-loop formalism in which state estimation and control are not designed separately. In the form used here, the free-energy functional is simply a weighted sum of squared prediction errors: each sensory channel’s mismatch between observed and predicted signals is squared, multiplied by that channel’s precision, and summed, together with a penalty on the belief’s rate of change (Section 3.3); minimizing it moves the belief toward the precision-weighted sensory evidence and moves action toward the preferred sensory state. This is the property exploited here: changes in sensory precision alter belief updating and then propagate to postural action through the shared perception–action loop. Crucially, prediction errors are precision weighted. Precision, the inverse variance or confidence assigned to a sensory channel, determines how strongly that channel contributes to belief updating and therefore to action (Feldman and Friston, 2010; Parr et al., 2022). This suggests a minimal mechanistic hypothesis: postural sensory reweighting is context-dependent precision control over vestibular, proprioceptive, and visual prediction errors.

The present study tests that hypothesis in a deliberately minimal model. We do not attempt a complete biophysical model of the vestibulospinal tract, a clinical account of vestibular disorders, or a full musculoskeletal simulation. Instead, we target the visual-scene and support-surface perturbation conditions that underlie the posturography benchmark for sensory reweighting: visual and proprioceptive conflicts are imposed while vestibular observations provide the body-state reference. We ask whether the core signature of sensory reweighting can be generated in a one-link quietstanding body model with a single ankle-torque action variable. This restricted model is useful because it isolates the mechanism: the body dynamics, action optimizer, and restorative sensory goal (the preferred sensory trajectory toward upright stance) are held fixed while only the imposed sensory precisions change.

The contribution is threefold. First, we formulate quiet-standing sensory reweighting as a minimal continuous-time mathematical model that integrates vestibular, proprioceptive, and visual observations, with intrinsic ankle stiffness and damping included in the body model. Second, we derive the model’s steady-state behavior in closed form: the perturbation-induced belief bias, a gradient-based reweighting index (the per-channel state-update contribution), the steady posture shift, and the posture-shift reduction all follow from the sensory precisions, the goal dynamics, and the intrinsic stiffness, together with a linear stability condition. These predictions are calculated analytically from the specified model parameters. The parameters are not fitted to the simulation results. Third, we test these predictions in 115 closed-loop trials that vary the target precision over a 32-fold range, scale all precisions together, add intrinsic stiffness up to the gravitational load stiffness, change the goal dynamics across the stability boundary, and change body mass and height; we also show the time course of belief, gradient, goal, and torque when the precision of one channel is switched within a trial. Channel-selective precision reduction reproduces Peterka-style flattening of perturbation-response curves, intrinsic stiffness bounds that flattening for physical support tilts, and global precision reduction does not flatten the curves. We state explicitly where the predictions hold and where the loop is unstable or unsettled.

## 2 Related Work

### 2.1 Sensory reweighting in human postural control

Human stance control has long been understood as a multisensory problem rather than a purely spinal or fixed-gain feedback loop. Visual, vestibular, and somatosensory signals each provide partial information about body orientation, and their reliability changes with lighting, support-surface motion, fatigue, and task context. Early dynamic-posturography work showed that stance control adapts to altered support and visual conditions, including in patients with vestibular deficits (Nashner et al., 1982). The empirical phenomenon most directly targeted here is sensory reweighting: the nervous system changes the effective contribution of sensory signals as their reliability changes. Peterka’s system-identification experiments showed that human postural responses to visual and support-surface rotations change with perturbation amplitude and sensory context, motivating perturbation-response curves and sensory weights as core descriptors of quiet-standing control (Peterka, 2002). Related experiments showed simultaneous reweighting of visual and tactile information during posture control (Oie et al., 2002). Subsequent work emphasized that these weights are dynamically regulated rather than fixed constants (Peterka and Loughlin, 2004), and later studies explicitly modeled and measured sensory-reweighting dynamics (Carver et al., 2006; Assländer and Peterka, 2014). Mergner’s review similarly framed reactive stance control as an interaction between body mechanics, sensory transformation, and context-dependent feedback organization (Mergner, 2010).

Work since 2015 has sharpened the empirical picture in three ways that bear on the present model. First, the dynamics of reweighting have been measured after cue removal or addition, with results depending on the sensory condition and response measure. Following visual or proprioceptive cue transitions, stimulus-evoked sway and sway variability can have different transition dynamics; faster adjustment after cue removal occurs in some, but not all, conditions (Assländer and Peterka, 2016). In a tandem-stance task, withdrawal of visual or haptic information produced faster adjustment of sway than its addition (Honeine et al., 2015). Second, system-identification studies have quantified channel weights and their dependence on perturbation amplitude, age, and disease: proprioceptive weight falls with disturbance amplitude and rises with age (Pasma et al., 2015); model-estimated vestibular weight falls with the severity of vestibular dysfunction (van Kordelaar et al., 2018); and a clinical implementation of the independent-channel model provides normative weights and feedback delays (Peterka et al., 2018). Third, a distinct form of context-dependent sensory modulation is the anticipatory reduction of vestibular-evoked balance responses before transitions between standing and walking (Tisserand et al., 2018). This anticipatory modulation does not establish the same mechanism as reliability-dependent sensory reweighting. Together, these studies motivate explicitly representing sensory contributions and their context dependence, while distinguishing the tasks and mechanisms involved.

The present model does not fit human posturography data quantitatively. Instead, it uses Peterka-style curve flattening as a qualitative benchmark: when the model treats a sensory channel as unreliable, the perturbation-response curve should flatten without changing the body dynamics or manually retuning a feedback controller.

### 2.2 Computational models of multisensory stance control

Several computational accounts have formalized postural control as state estimation and feedback control under uncertainty. Adaptive sensory-integration models of stance control showed how sensory channel gains can be adjusted as environmental conditions change (van der Kooij et al., 2001). Stochastic and multisensory fusion models further related postural sway to the integration of noisy sensory streams (Kiemel et al., 2002). Work on the informational content of postural signals also emphasized that velocity-related sensory information can be more useful than position or acceleration alone for controlling upright stance (Jeka et al., 2004). These models are important because they moved the field beyond single-channel reflex explanations and toward closed-loop estimationcontrol accounts. More recent modeling has identified the plant and the neural feedback separately in closed loop, showing why a change in body response need not by itself identify a change in neural feedback (Hwang et al., 2016). A review distinguished immediate postural adjustments from longer-term changes with aging, learning, and rehabilitation (Chiba et al., 2016). Modular disturbance-estimation-and-compensation control has also been implemented on a 14-degree-of-freedom humanoid, with coordination emerging without a central full-body controller (Lippi and Mergner, 2017). In a one-degree-of-freedom robotic balancing task, non-predictive, continuous predictive, and intermittent predictive models were compared; the intermittent predictive model better reproduced the mean frequency, sample entropy, and response features of the nonlinear 0.2–2 Hz control component (Loram et al., 2022). For the body model, simulations support a single inverted pendulum as a useful approximation to center-of-mass control during quiet standing on a large, rigid support with sufficient ankle torque, despite ankle–hip motion (Morasso et al., 2019). A delayed feedback model analyzed coadaptation of intrinsic ankle stiffness, feedback gain, and delay, with higher stiffness or delay requiring lower gain within that model’s assumptions (Le Mouël and Brette, 2019). Experiments with robot-imposed sensorimotor delays found reduced vestibular-evoked contributions before training and partial restoration with learning (Rasman et al., 2021).

Bayesian and optimal-control approaches provide a broader normative background (Todorov and Jordan, 2002). In sensorimotor learning, Bayesian models predict that uncertain sensory signals should be weighted less than reliable sensory signals (Ernst and Banks, 2002; Körding and Wolpert, 2004; Knill and Pouget, 2004). In a full-body task, Stevenson and colleagues showed that visual uncertainty could be incorporated into models combining Bayesian estimation and nonlinear feedback control (Stevenson et al., 2009). These studies motivate the present focus on reliabilitydependent weighting, but they generally separate state estimation and control policy design. Our contribution is to express the reweighting mechanism inside a single variational objective, and to measure the actual contribution of each sensory channel to the belief update.

### 2.3 Active inference, precision, and motor control

Active inference treats perception and action as coupled minimization of variational free energy (Friston, 2010; Friston et al., 2010). In motor control, Adams, Shipp, and Friston argued that descending motor signals can be understood as predictions of proprioceptive sensations rather than classical commands (Adams et al., 2013). This view is especially relevant to posture because reflex-like action can be interpreted as the fulfillment of predicted sensory states. In the free-energy account of attention, precision represents the inverse uncertainty assigned to prediction errors and modulates the gain of these error signals during inference (Feldman and Friston, 2010).

A self-contained mathematical review describes the continuous-state formulation of active inference (Buckley et al., 2017). Under a linear generative model and specific approximation assumptions, active inference can yield proportional-integral-derivative control, with precision parameters related to controller gains (Baltieri and Buckley, 2019). In robotics, continuous active inference has controlled a simulated PR2 arm under sensory noise (Pio-Lopez et al., 2016) and supported reaching and head tracking on a real iCub humanoid under sensory noise and sensorimotor conflict (Oliver et al., 2022). A review of predictive processing in cognitive robotics identified precision weighting as a direction for further research and noted limited attention to the temporal dynamics of prediction errors (Ciria et al., 2021). In a hand-movement task under visuo-proprioceptive conflict, an active-inference simulation that increased the expected precision of the instructed modality reproduced aspects of human task performance (Limanowski and Friston, 2020). Synthetic perturbations of precision and other generative-model parameters also reproduced specific reflex and coordination abnormalities in simulated upper-limb examinations (Parr et al., 2021). Precision has been proposed to operate through synaptic gain control in predictive-processing accounts of attention (Kanai et al., 2015); a review of body representation similarly considered top-down precision control as an account of flexible integration of visual and proprioceptive feedback (Limanowski, 2022). These studies provide computational and conceptual precedents, rather than empirical validation of the present stance model.

Recent surveys have positioned active inference as a framework for robotic state estimation and control under uncertainty, while also noting challenges in scaling, tuning, and benchmarking (Lanillos et al., 2021). A complementary survey has focused specifically on continuous-time active inference models of motor control (Priorelli et al., 2024). The present work aligns with that literature but deliberately narrows: it tests one mechanistic claim, namely that changing relative sensory precision is sufficient to generate closed-loop postural reweighting in a minimal stance model, and it derives the consequences in closed form.

### 2.4 Gap addressed by the present work

Prior postural-control models have characterized sensory reweighting, and Bayesian/optimal-control models have explained why uncertain sensory signals should be weighted less. Active inference provides the additional link that precision weighting affects both belief updating and action within a single objective. What has been missing is a minimal closed-loop demonstration showing that channel-specific precision changes reduce the influence of unreliable sensory perturbations on belief updating and, in turn, on perturbation-driven posture, together with an analytical account of how those reductions follow from the precision parameters. This is the gap addressed here.

## 3 Methods

### 3.1 One-link body model

The body is modeled as a sagittal-plane one-link inverted pendulum rotating about the ankle (Winter et al., 1998); its geometry is shown in Fig. 1. This model simplifies body mechanics to examine how sensory precision affects postural control. The one-link approximation excludes multi-joint postural strategies.

**Figure 1.**
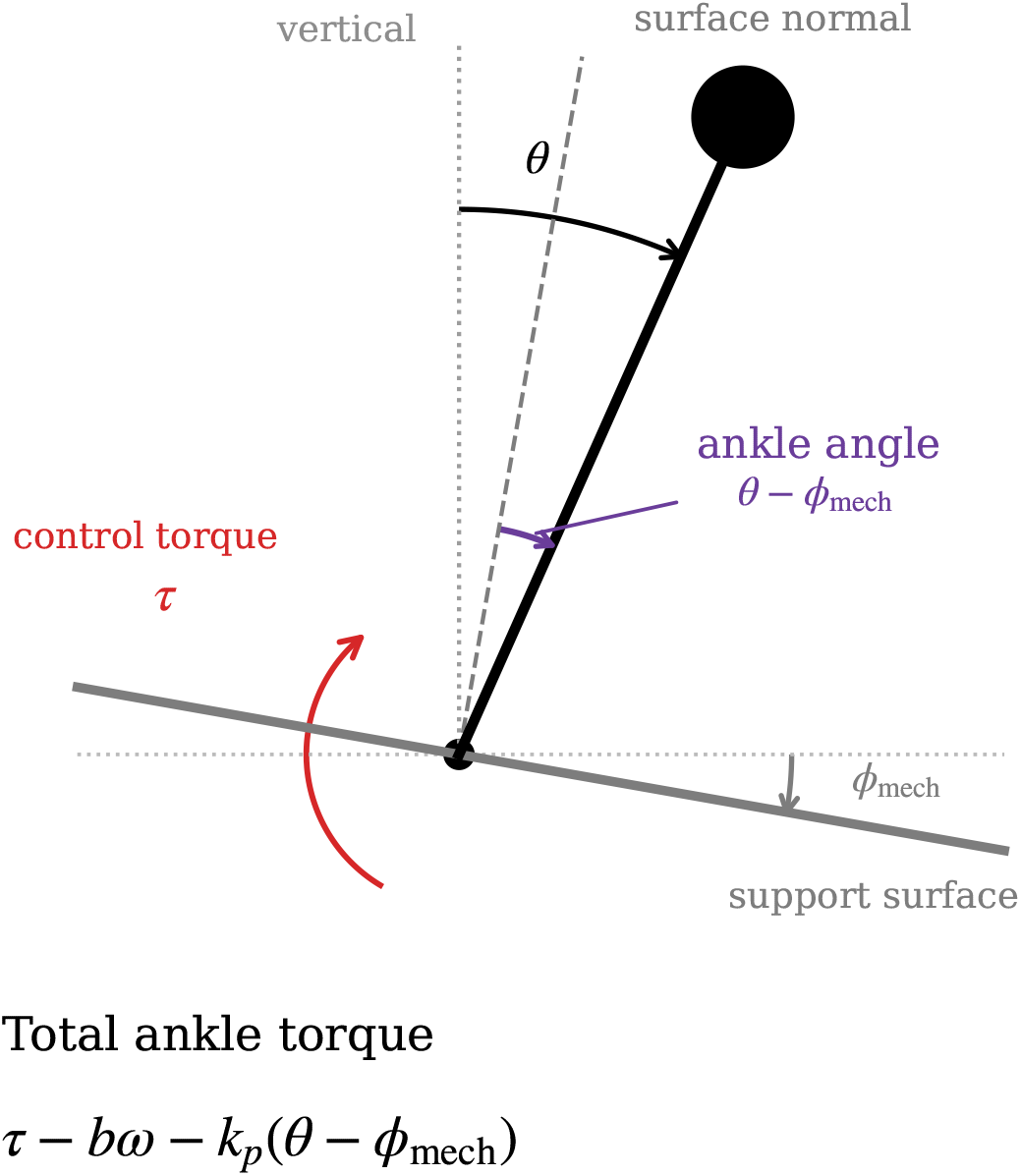
Physical geometry and ankle torques. The body angle *θ* is measured from vertical, and *ϕ*_mech_ is the physical support-surface angle. Their difference is the actual ankle angle. The control torque *τ*, damping torque −*bω*, and intrinsic spring torque −*k*_*p*_(*θ*−*ϕ*_mech_) sum to the total modeled ankle torque. Positive rotations are clockwise.

The body state is *x* = [*θ, ω*]^*⊤*^, where *θ* is body tilt about the ankle and 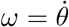 is the corresponding angular velocity. The action variable *τ* is an idealized ankle-torque input selected by the A-step. The continuous-time body dynamics are

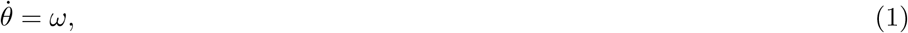

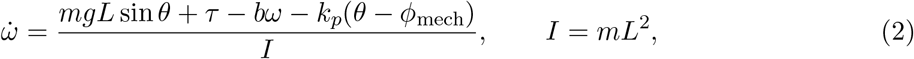

where *m* is the lumped body mass and *L* is the distance from the ankle to the center of mass, equal to its height above the ankle when the body is upright. The moment of inertia *I* = *mL*^2^ uses a point-mass approximation. The coefficient *b* represents passive ankle damping acting on the body angular velocity *ω*, and *k*_*p*_ is an effective intrinsic ankle stiffness. The support surface has mechanical tilt *ϕ*_mech_, so the modeled ankle angle is *θ*−*ϕ*_mech_. We represent intrinsic stiffness by the linear spring torque −*k*_*p*_(*θ* −*ϕ*_mech_), choosing zero ankle angle as its zero-torque reference. Sensory perturbations and their relation to physical support tilt are defined separately in Section 3.2.

In humans, deformation of muscle fibers, tendons, and other tissues spanning the ankle can change ankle torque without a change in neural drive, including when the muscles are already active (Sakanaka et al., 2018). Neural modulation of muscle force provides a further means of changing ankle torque, represented here by the control input *τ* selected by the A-step. These contributions involve the same muscle–tendon structures, and intrinsic stiffness can itself change with muscle activity and tendon loading (Sakanaka et al., 2018). Holding *k*_*p*_ constant within each condition, rather than allowing it to vary with *τ*, is therefore a modeling simplification. The model does not resolve neural circuitry, muscle activation, or tendon dynamics. The total modeled ankle torque is *τ* − *bω* − *k*_*p*_(*θ* − *ϕ*_mech_).

Near upright, the gravitational torque has a destabilizing slope of *mgL*, opposed by the intrinsic restoring stiffness *k*_*p*_. Experimental studies report intrinsic stiffness relative to this gravitational torque slope using the dimensionless ratio *k*_*p*_*/*(*mgL*), called relative or normalized stiffness and often expressed as a percentage (Loram and Lakie, 2002; Sakanaka et al., 2021). This normalization allows comparison across participants with different body masses and center-of-mass heights.

Loram and Lakie (2002) reported combined intrinsic ankle stiffness of *k*_*p*_*/*(*mgL*) = 0.91 ± 0.23 (mean ± SD) during quiet standing. Subsequent studies show that intrinsic stiffness varies with perturbation amplitude, ankle torque, tendon stretch, and participant (Sakanaka et al., 2018, 2021). In the normal-standing condition of Sakanaka et al. (2021), stiffness measured with 0.2° perturbations was 0.941 ± 0.216 *mgL* (mean ± SD), compared with 0.579 ± 0.123 *mgL* for 0.9° perturbations. Individual short-range values across the study ranged from 0.40 to 1.34 *mgL*. Thus measured intrinsic stiffness can exceed *mgL*, but its magnitude depends on the participant and measurement conditions; an estimate above *mgL* at small perturbations does not establish that stiffness remains above this threshold over larger movements.

The baseline uses *k*_*p*_ = 0 to isolate active restoring torque. Section 5.8 tests *k*_*p*_*/*(*mgL*) = 0, 0.5, 0.9, and 1.0 to assess its effects on the model predictions. These values are representative model settings, not bounds on measured human stiffness. With *k*_*p*_ = 0, support-surface tilt has no mechanical effect on the body, so the baseline support-tilt conditions are purely sensory perturbations. The sensory-only and physical support-tilt conditions (Section 3.2) differ mechanically only when *k*_*p*_ *>* 0.

The damping torque vanishes at equilibrium, so *b* affects the transient response but does not enter the steady-state predictions (Appendix A.3); its effects are tested in Section 5.8. The support surface is static between imposed tilt changes. Torques proportional to support-surface angular velocity are not included.

The body model was integrated with a time step of 0.005 s (0.0025 s in the convergence tests). Unless otherwise noted, simulations started from *θ*_0_ = 0.05 rad and 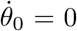. Angles are expressed in radians unless otherwise stated.

### 3.2 Multisensory observations

The model constructs vestibular, proprioceptive, and visual observations from the body state and the imposed sensory perturbations. Figure 2 shows this observation generation before the belief update. In the active-inference formulation used here, generalized observations collect an observation vector and its temporal derivatives. Each observation channel is represented with two temporal orders. The current observation vector, corresponding to the order-0 term, is

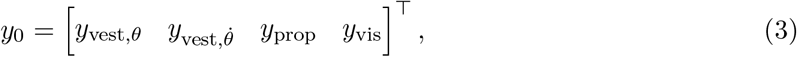

with channel values generated from the body state and perturbation variables as

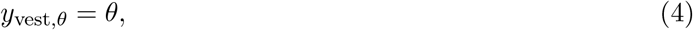

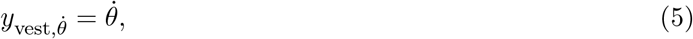

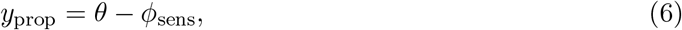

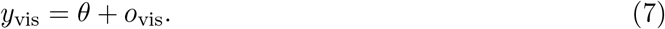

**Figure 2.**
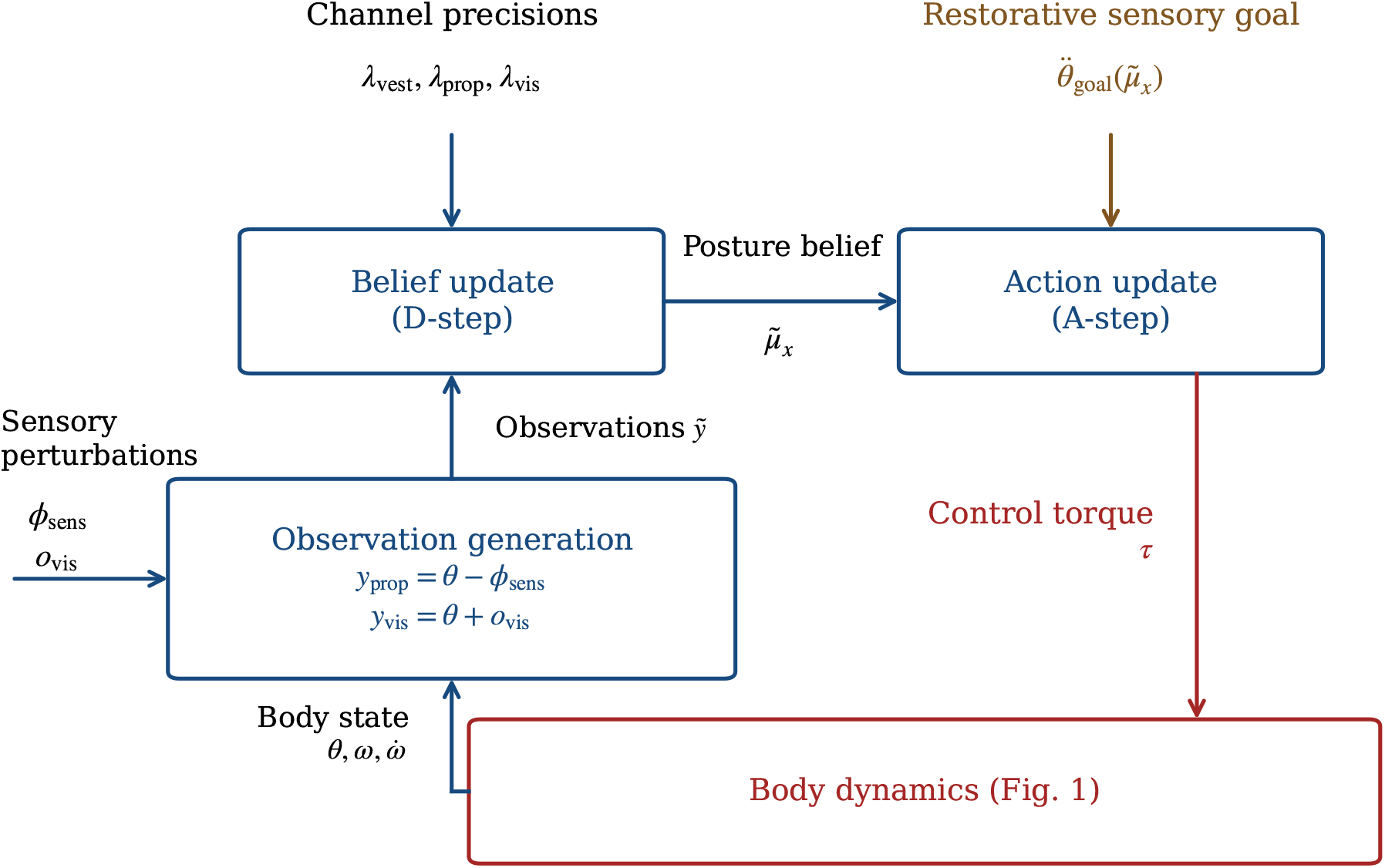
Observation construction and the perception–action loop. Body state and sensory perturbations are used to construct the observations supplied to the D-step. The displayed equations show the perturbed angle channels; vestibular observations and temporal derivatives are defined in Section 3.2. The D-step uses 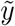 to update the posture belief through precision-weighted prediction errors; *ϕ*_sens_ and *o*_vis_ are not given to its prediction model as known offsets. The belief determines the restorative sensory goal and enters the A-step, which selects the control torque *τ*. The torque acts through the body dynamics of Fig. 1. Both updates minimize the shared precision-weighted free-energy form.

The first-order generalized-observation block is the time derivative of the current observation vector,

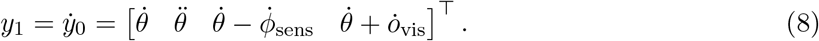

Generalized observations are then 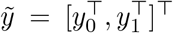. Throughout the model, a tilde denotes this temporal embedding of a variable and its derivatives.

Here, *ϕ*_sens_ is the support tilt used as the reference in the proprioceptive observation, and *o*_vis_ is the visual offset. These parameters are used to construct the observations supplied to the D-step; they are not supplied as known offsets to its prediction model 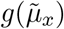 (Section 3.4). Because the actual ankle angle is *θ* − *ϕ*_mech_, the proprioceptive observation can also be written as

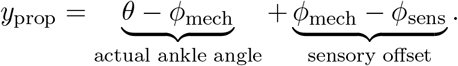

Thus the sensory offset relative to the actual ankle angle is *ϕ*_mech_ − *ϕ*_sens_, not *ϕ*_sens_ alone. A nonzero physical tilt produces no such offset when *ϕ*_sens_ = *ϕ*_mech_; a sensory-only perturbation with *ϕ*_mech_ = 0 produces offset −*ϕ*_sens_. For a tilt perturbation of amplitude *ϕ*, the sensory-only condition sets *ϕ*_sens_ = *ϕ* and *ϕ*_mech_ = 0, whereas the physical condition sets *ϕ*_sens_ = *ϕ*_mech_ = *ϕ*. The two coincide exactly when *k*_*p*_ = 0. In the main simulations, these observation equations were deterministic: sensory conflict was introduced by the visual offset or support-surface tilt terms, not by adding random sensor noise. This choice does not assume that vestibular sensing is biologically noise-free. Rather, the present perturbation conditions model visual-scene and support-surface perturbations, for which vestibular observations serve as the body-state reference against which visual and proprioceptive conflicts are defined. Direct vestibular bias would define a different perturbation condition, such as galvanic vestibular stimulation, and was left outside the present minimal analysis.

In the simulations, perturbation amplitudes were constant during the perturbation window, so 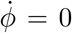 and 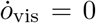 away from perturbation onset and offset. For this temporal embedding, the time derivative of the vestibular angular-velocity observation was evaluated from the simulated body dynamics. This treatment provides the derivative term required by the embedding and should not be interpreted as assuming that the vestibular periphery directly measures angular acceleration; in biological postural control, acceleration-like quantities would be implicit in central state estimation and sensorimotor dynamics.

### 3.3 Continuous active inference model

Fig. 2 summarizes the perception-action loop used in the model. Standing control is implemented as continuous active inference: within each body-dynamics integration step the model alternates between calculation of a belief update and calculation of an action update, in that order, and the resulting torque is then applied to the body model for that step; both updates reduce the same precision-weighted free-energy form defined below. The belief update, called the D-step, changes the posture belief for the current sensory observation; it follows the belief-update step of dynamic expectation maximization (DEM) over temporally embedded states (Friston et al., 2008). The action update, called the A-step, changes ankle torque toward the restorative sensory goal through an action-dependent observation mapping; it extends the same objective to action in the manner of continuous active inference (Friston et al., 2010). This subsection defines the belief variables and the shared objective; the two subsections that follow specify the two updates.

The two-order generalized state corresponding to the body state *x* = [*θ, ω*]^*⊤*^ is 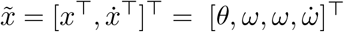. The corresponding generalized-state belief is

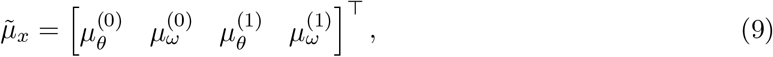

where superscripts denote temporal order in the embedding. Thus, 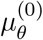 and 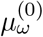 are the order-0 beliefs about body tilt and angular velocity, and 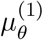 and 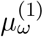 are the first-order embedding components. The belief update treats 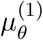 and 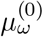 as distinct coordinates: they estimate the same physical quantity but are not constrained to coincide, because they enter the objective through different terms. The component 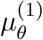 enters the smoothness term and the first-order angle-channel predictions, whereas 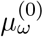 enters only the order-0 angular-velocity prediction.

The active-inference standing model uses the same precision-weighted free-energy form for belief updating and action selection. Let 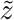 denote the generalized observation vector used for comparison, and let 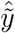 denote the corresponding predicted generalized observation. The shared form is

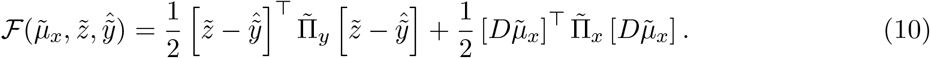

Here, *D* is the temporal shift operator for the generalized-state embedding, and 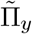 and 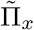 are generalized sensory and state precisions. The three arguments of *ℱ* 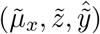 are the belief, the sensory reference, and the predicted observation; which of them varies depends on the update. In the D-step, 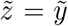 is the current observation and is held fixed, whereas 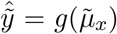 depends on the belief, so the gradient with respect to 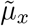 passes through *g* (Section 3.4). In the A-step, the belief and the goal vector are held fixed and 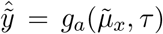 depends on the torque, so the gradient is taken with respect to *τ* through *g*_*a*_ (Section 3.5). The first term penalizes the precision-weighted mismatch 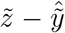. In the D-step, 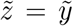, so this mismatch is the sensory prediction error for the current observation. In the A-step, 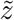 is the A-step goal vector defined below, so the same mismatch is the error relative to the restorative sensory goal. The second term is a smoothness penalty on the belief trajectory: with the truncated two-order embedding used here,

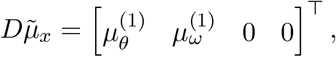

because second-order belief components are not represented. Thus, this term penalizes the first-order belief components and leaves the order-0 posture belief unconstrained. In the standard DEM formulation, the second term is the generalized state-prediction error 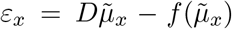 for a dynamics prior *f* (Friston et al., 2008); the present implementation uses *f ≡* 0, so *ε*_*x*_ reduces to the smoothness penalty above and no explicit inverted-pendulum dynamics prior enters the belief update. This is a deliberate modeling restriction that isolates the contribution of sensory precision weighting.

### 3.4 D-step belief update

The observation model maps the generalized-state belief to the predicted generalized observation 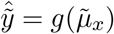. Following the channel order in 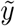, the observation model is

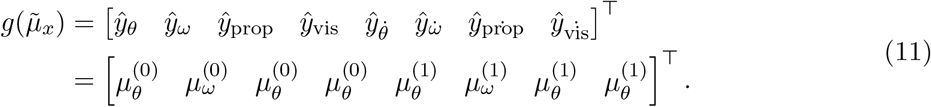

Thus, angle-level channels are predicted from the tilt belief 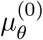 and its first-order component 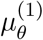, whereas the vestibular angular-velocity channel is predicted from 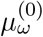 and its derivative component 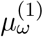. Repeated entries in 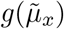 denote the same belief value reused across different sensory channels. For example, the three occurrences of 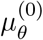 predict body tilt, proprioceptive angle, and visual angle from the same body-tilt belief; channel-specific discrepancies arise in the actual observation vector 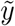 through support-surface and visual perturbations.

External visual and support-surface perturbations enter the actual observation 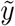, not the prediction 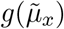, thereby producing sensory prediction errors. Substituting the current observation and this observation model into the shared free-energy form gives the D-step objective:

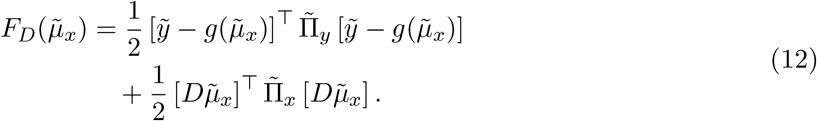

In the reported simulations, the D-step uses the gradient-descent mode

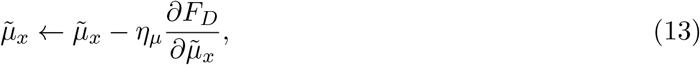

with D-step optimizer step size *η*_*µ*_ = 0.001 and, at the representative setting, 32 D-step iterations per body-dynamics integration step. The convergence tests also used 16 and 64 iterations. Here, 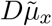 enters the update only through the smoothness term of *F*_*D*_; it is not added as a separate drift term in the update equation, as it would be in a full generalized-filtering scheme (Friston et al., 2008). The D-step is therefore an iterative gradient optimization of the current precision-weighted prediction errors. This choice does not move the equilibria analyzed below: at the late-window fixed point the first-order belief components vanish, so 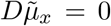 and the gradient-descent fixed point is also a stationary point of the generalized-filtering flow.

In the D-step, posture is estimated from sensory prediction errors without using the inverted-pendulum equation as an internal dynamics prior for belief updating. A linearized inverted-pendulum prior was also tested but destabilized the belief-driven baseline because the open-loop unstable prior contaminated state estimation; it was not used in the reported results.

### 3.5 A-step action update and the restorative sensory goal

The A-step requires a preferred sensory trajectory, which we call the restorative sensory goal. In traditional control-system models of stance, a set point (typically zero body-sway angle) defines an error signal from which a neural controller computes a corrective ankle torque. The present model also has a set point of zero tilt, but it is expressed at the level of a preferred sensory signal rather than as an angle error: the preferred vestibular angular acceleration is a restoring acceleration computed from the current tilt and velocity beliefs,

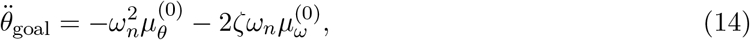

where 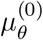and 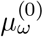 are the order-0 D-step beliefs about body tilt and angular velocity, *ω*_*n*_ is the natural frequency of the restorative goal, and *ζ*is its damping ratio. The representative values are *ω*_*n*_ = 4.0 rad/s and *ζ*= 1.0 (Table 1); Section 5 also tests other goal frequencies and damping ratios. The preferred acceleration is therefore not zero in general; it is zero only when the believed tilt and velocity satisfy 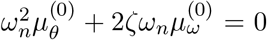, and for a stationary belief 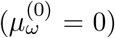 this requires 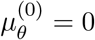: the implicit set point is zero believed tilt. Because the belief is biased by sensory conflict (Section 4), the actual body tilt at equilibrium is not zero. A static goal with zero preferred acceleration was tested as an alternative and did not stabilize standing, because it provides no tilt-dependent restoring effect. Although Eq. (14) has the form of a second-order return-to-upright acceleration law, it is not a separately applied proportional-derivative controller: the goal enters the model only as the preferred value of a sensory channel, and torque is still selected by minimizing the shared free-energy form. Its correspondence to stiffness and damping gains is made explicit below. The A-step sensory goal vector is

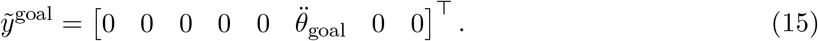

**Table 1:** Model, optimizer, and protocol parameters. “Representative” values define the baseline setting; “Explored” lists additional values tested in the simulation blocks of Section 5. Dashes indicate fixed values. The combined-conflict condition uses a +5° visual offset and a −5° sensoryonly support tilt. Hold and trial durations follow the slow-mode rule of Section 3.8, rather than being independent test factors. The preliminary checks and precision-switch trials use the schedules specified in Sections 5.1–5.2 and 5.9, respectively.

| Parameter | Representative | Explored | Meaning |
|---|---|---|---|
| *Body mechanics* |  |  |  |
| *m* | 70.0 kg | 50, 90 | body mass |
| *L* | 1.0 m | 0.8, 1.2 | center-of-mass height |
| *I* = *mL*<sup>2</sup> | 70.0 kg m<sup>2</sup> | follows *m*, *L* | body moment of inertia |
| *b* | 2.0 N m s rad<sup>−1</sup> | 0, 20 | passive damping |
| *k*<sub>*p*</sub> | 0 | 0.5, 0.9, 1.0 *mgL* | intrinsic ankle stiffness |
| *g* | 9.81 m s<sup>−2</sup> | – | gravitational acceleration |
| *Sensory model and goal* |  |  |  |
| *ω*<sub>*n*</sub> | 4.0 rad s<sup>−1</sup> | 3.0, 3.5, 5.0 | goal natural frequency |
| *ζ* | 1.0 | 0.5, 1.5 | goal damping ratio |
| *λ*<sub>vest</sub>, *λ*<sub>prop</sub>, *λ*<sub>vis</sub> | 8, 8, 8 | target 0.5–16 | channel precisions |
|  |  | all ×0.25, ×4; (1, 1, 1) |  |
| Π̃<sub>*x*</sub> | 2.0*I*<sub>4</sub> | – | state smoothness precision |
| *s*<sub>*y*</sub> | 1.0 | – | sensory fluctuation smoothness |
| *Action constraints* |  |  |  |
| Δ*τ*<sub>max</sub> | 150 N m | – | torque-increment bound |
| *τ*<sub>max</sub> | 300 N m | – | torque bound |
| *Numerical settings* |  |  |  |
| *η*<sub>*μ*</sub> | 0.001 | – | D-step optimizer step size |
| D-step iterations | 32 | 16, 64 | belief iterations per integration step |
| A-step iterations | 1 | – | Newton iterations per step |
| *δ* | 10<sup>−6</sup> | – | Newton regularization constant |
| Integration step size | 0.005 s | 0.0025 s | body-dynamics step |
| *Simulation protocol* |  |  |  |
| Perturbation amplitude | +5° | 0, ±2, ±4, ±8° | visual offset or support tilt |
|  |  | +0.25° | near stability boundary |
| Perturbation onset | 2 s | – | see Section 5 |
| Perturbation hold | 14 s | up to 26 s (rule) | max(14 s, 10 *τ*<sub>slow</sub>), cap 60 s |
| Trial duration | 18 s | up to 30 s (rule) | hold + 4 s |

Action changes predicted observations through a separate physical action-observation mapping,

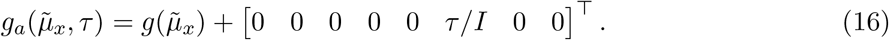

Here, *I* is the same body-model inertia parameter as in Eq. (2). In the biological interpretation, the term *τ/I* represents an effective calibrated action-to-acceleration mapping rather than explicit knowledge of the body’s moment of inertia. In Eq. (16), ankle torque affects the predicted vestibular angular-acceleration channel only. This action-observation mapping imposes the physical constraint that action cannot change posture observations instantaneously: torque acts first on angular acceleration through the body dynamics, and angle-level observations change only after subsequent body-model integration.

Evaluating the shared free-energy form with the goal vector and the action-dependent prediction gives the A-step objective:

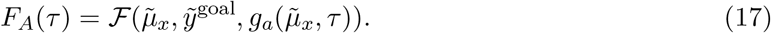

*F*_*A*_ is an instantaneous variational objective in which a preferred observation replaces the current observation; it is not an expected-free-energy evaluation over future policies. During this update, 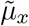 held fixed, so the smoothness term in Eq. (10) is constant with respect to *τ* and does not contribute to the action gradient. For the vestibular angular-acceleration channel, the A-step prediction error is

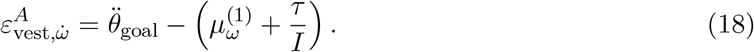

Eq. (14) does not overwrite the belief acceleration 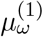 in 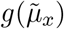; instead, it supplies the desired value in the goal observation. The A-step uses a scalar Newton step,

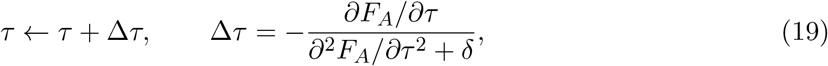

where *δ* is a small regularization constant in the Newton denominator; the increment is clipped to |Δ*τ*| *≤Δτ*_max_ and the torque to |*τ*| *≤ τ*_max_. This Newton update scales the torque increment by the local curvature of *F*_*A*_, rather than by a separately tuned action learning rate. Throughout the paper, the term action optimizer refers to this fixed A-step scheme: the objective *F*_*A*_ together with the Newton update, its regularization, and its bounds.

#### Correspondence to stiffness and damping gains

For comparison with control-system models that express the neural controller as a stiffness and a damping factor, the torque that one Newton step selects outside the bounds and with *δ →* 0 is 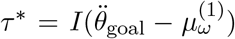 (Section 4). Substituting Eq. (14) gives

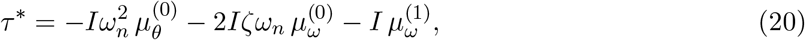

so the goal law acts on the believed tilt and velocity with a stiffness factor 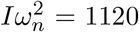 N m rad^−1^ and a damping factor 2*Iζω*_*n*_ = 560 N m s rad^−1^ for the parameters in Table 1, plus a third term proportional to the believed angular acceleration 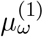 that has no counterpart in a proportionalderivative controller and acts as an added effective inertia (Appendix A.3). The scaling from the normalized goal acceleration to a physical torque happens in the action-observation mapping of Eq. (16), whose factor *τ/I* makes the Newton step solve for torque in N m. The stiffness factor exceeds the gravitational load stiffness *mgL* = 687 N m rad^−1^, leaving a net closed-loop restoring stiffness of 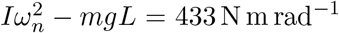 at *k*_*p*_ = 0 (+*k*_*p*_ when intrinsic stiffness is present); this is the linear stability condition derived in Section 4. These factors are properties of the restorative goal, not of the ankle’s mechanical properties (*b, k*_*p*_), and the correspondence is a mathematical equivalence of the closed loop, not a claim that active inference outperforms, or is implemented differently from, a feedback controller with the same gains.

### 3.6 Channel-block sensory precision

Sensory precision is assigned per channel:

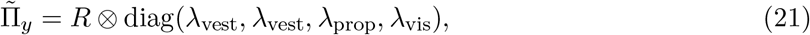

where *R* is a fixed temporal-embedding precision structure, not a quantity fitted from data. It is constructed from the standard DEM smooth-fluctuation assumption (Friston et al., 2008), in which a smoothness parameter *s*_*y*_ sets the assumed temporal correlation width of sensory fluctuations. For the two-order embedding used here, this construction gives 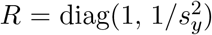, so *s*_*y*_ only rescales the weight of the first-order prediction errors relative to the order-0 ones; with *s*_*y*_ = 1.0, *R* reduces to *I*_2_. The sensory precision is then the diagonal matrix

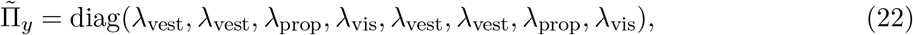

whose entries follow the same channel order as 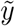 and 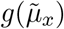: each channel’s precision weights both its order-0 and its first-order prediction error. The scalar precisions *λ*_vest_, *λ*_prop_, and *λ*_vis_ weight vestibular, proprioceptive, and visual prediction errors, respectively. In this minimal model, the vestibular angle and angular-velocity components are treated as one modality block and therefore share *λ*_vest_; this is not a biological claim that their reliabilities must be identical. Precision triples below are reported as (*λ*_vest_, *λ*_prop_, *λ*_vis_). Equal precision was (8, 8, 8) for vestibular, proprioceptive, and visual channels. To model an unreliable sensory channel, the target channel precision was reduced to 1, yielding (8, 8, 1) for visual precision reduction and (8, 1, 8) for proprioceptive precision reduction. The values 1 and 8 are dimensionless model precisions, not biological bounds. They were chosen to provide a fixed high–low precision contrast around the stable equal-precision baseline while keeping the body dynamics, goal, and action optimizer unchanged across conditions. The channel precisions were experimenter-specified condition parameters, not variables inferred from sensory data or learned online. In the steady-state analyses, each simulation used the precision values specified by its condition, held fixed for the whole trial; the target-channel precision *λ*_*t*_ was imposed directly at discrete values between 0.5 and 16 (with the two non-target precisions at 8), without any intermediate reliability variable. In the time-course analysis (Section 5.9), the target precision was switched within a trial, from 8 to 1 at 7 s and back to 8 at 12 s, while the belief, torque, and body state were carried across the switch; from the switch step onward both the belief update and the action update use the new precision. The model therefore tests whether imposed precision changes are sufficient to generate sensory reweighting; it does not explain how the human nervous system estimates or updates those precisions, and it does not predict the time course of such estimation.

Operationally, these precisions are the model’s sensory weights. They do not act as a separate normalized weighting rule; they enter the free-energy objective through 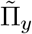. Here, precision denotes a model parameter that weights sensory prediction errors; it is not an empirical estimate of physical sensor-noise variance. Under a Gaussian observation likelihood this parameter corresponds to an inverse observation variance, and Appendix A.7 tests this correspondence directly by pairing each precision with a matched observation-noise variance 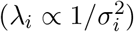. Consequently, writing *F*_*y*_ for the sensory term of *F*_*D*_, the D-step gradient can be read, up to sign, as a sum of channel contributions,

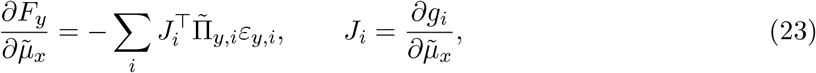

where *i* indexes the vestibular, proprioceptive, and visual channel blocks, each block collecting a channel’s components at both temporal orders. Here, *ε*_*y,i*_ is block *i* of the sensory prediction error 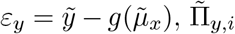 is the corresponding diagonal block of 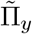, and *g*_*i*_ collects the components of *g* in block *i*. Reducing *λ*_*i*_ therefore reduces how strongly the prediction error from channel *i* pulls the posture belief during the D-step. The resulting belief then enters the restorative sensory goal in Eq. (14) and the A-step action-observation mapping, so the altered sensory weighting propagates to ankle torque without changing the body dynamics or the action optimizer.

### 3.7 State-update gradient contribution

To quantify reweighting at the mechanism level, we measured how strongly each sensory channel would update the posture belief during the D-step. In active inference, sensory prediction errors update the state belief through the free-energy gradient with respect to 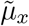. If the precision of a channel is reduced, the same sensory conflict should make a smaller contribution to that gradient. For channel *i*, the channel-masked sensory free energy is

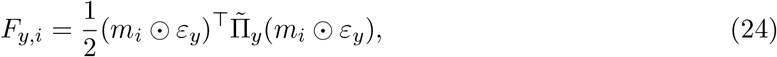

where *⊙* denotes the element-wise product and *m*_*i*_ is a binary mask vector that selects the vestibular, proprioceptive, or visual components of *ε*_*y*_ and sets the other sensory components to zero. The signed angle gradient of channel *i* is

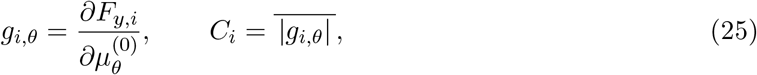

and the contribution index *C*_*i*_ is the mean of |*g*_*i,θ*_| over the evaluation window. *g*_*i,θ*_ has units of precision times radians; a gradient-descent belief update moves 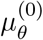 along −*g*_*i,θ*_, so its sign indicates the direction in which channel *i* pulls the tilt belief and its magnitude how strongly. *C*_*i*_ is therefore the absolute update pressure that channel *i* exerts on the estimated body tilt; *C*_vis_ and *C*_prop_ denote this quantity for the visual and proprioceptive channels. The angle-only definition avoids mixing quantities with different units during transients. At steady state the angular-velocity gradient components vanish for every channel, so *C*_*i*_ also equals the Euclidean norm of the gradient over the tilt and angular-velocity beliefs. Gradients were evaluated by automatic differentiation rather than finite differences. We used the absolute contribution *C*_*i*_ rather than each channel’s share of the total contribution, because a share can remain unchanged even when the channel’s absolute influence on belief updating decreases. In the simulation experiments, the reported state-update gradient contributions are values of *C*_*i*_, and the reported gradient reductions are condition-wise changes in *C*_*i*_, both evaluated for the perturbation-relevant channel: visual for visual-offset perturbations and proprioceptive for support-surface-tilt perturbations. *g*_*i,θ*_ is a model error-scale quantity, not a neural signal.

### 3.8 Simulation protocol and evaluation window

Table 1 groups the simulation parameters by function and lists the baseline values and the alternatives tested. Unless otherwise noted, simulations apply the perturbation at 2 s, hold it for 14 s (removal at 16 s in an 18 s trial), and evaluate the late response window as the mean posture over the final 2 s of the hold (14 s to 16 s). For conditions whose linearized slow time constant *τ*_slow_ is at least 1.5 s, the hold duration was set to max(14 s, 10 *τ*_slow_), subject to a cap of 60 s specified before the runs, and the window is the final 2 s of that longer hold (Appendix A.5). The precision-switch trials follow their own schedule (Section 5.9), and the two preliminary checks of Sections 5.1 and 5.2 use a shorter perturbation interval of 2 s to 6 s. The 5 s to 6 s evaluation window in that protocol still contains a closed-loop transient (Appendix A.5), so it is not used for steady-state verification. Long holds serve only to verify the steady-state theory; they are not claims about the speed of human postural responses.

## 4 Fixed-point analysis

Before turning to simulations, we characterize the model’s steady-state behavior and its linear stability in closed form. The analysis applies to the constant-perturbation regime used in the late response window, with beliefs equilibrated and the body dynamics linearized around upright; the corresponding nonlinear equilibrium is given in Appendix A.3 and is the quantity against which simulations are compared. Full derivations are given in Appendix A. Each result is a parameter-explicit prediction in terms of the precisions, the goal dynamics (*ω*_*n*_, *ζ*), the body parameters, and the intrinsic stiffness *k*_*p*_; the simulation experiments in Section 5 test these predictions in the full nonlinear closed loop and report where they fail.

### 4.1 Belief bias is set by relative precision

For a constant visual offset *o* and support-surface tilt *ϕ*, the order-0 D-step fixed point is the precision-weighted sensory average (Appendix A.1), so the perturbation-induced belief bias is

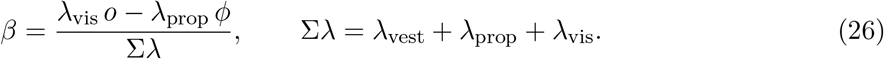

Two properties of Eq. (26) matter for what follows. First, *β* depends only on precision ratios: rescaling all precisions together, as in the global low-precision regime, leaves *β* unchanged. Second, the state smoothness precision 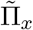 does not enter Eq. (26), because the smoothness term penalizes only the first-order belief components; the bias predictions are therefore not tuned by this free parameter.

**Table A.1:** Closed-form and simulated visual state-update contributions for imposed visual precision λ_vis_ (non-target precisions 8).

| λ<sub>vis</sub> | Closed form | Simulation |
|---|---|---|
| 1.00 | 0.0821 | 0.082 |
| 2.75 | 0.2048 | 0.205 |
| 4.50 | 0.3065 | 0.306 |
| 6.25 | 0.3922 | 0.392 |
| 8.00 | 0.4654 | 0.465 |

### 4.2 State-update contributions in closed form

At the fixed point, the state-update contribution of the perturbed channel has the closed form

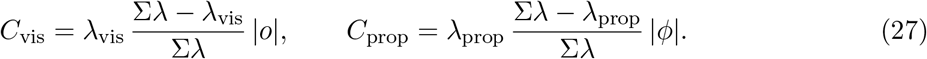

Once the sensory precisions and perturbation amplitude are specified, these expressions predict the state-update contribution *C*_*i*_ without additional parameter fitting. They apply to all precision conditions examined below, including graded target-channel precisions and the global low-precision regime. Intrinsic stiffness *k*_*p*_ does not enter these expressions because it does not enter the belief update.

When only one sensory channel is perturbed, its absolute contribution at the fixed point equals the sum of those of the unperturbed channels. The perturbed channel therefore accounts for half of the total absolute contribution, regardless of the sensory precisions (Appendix A.2). Precision changes can alter the absolute contributions while leaving this share unchanged. We therefore use absolute contributions to quantify reweighting.

### 4.3 Closed-loop posture bias and a stability condition

The A-step admits an exact solution (Appendix A.3): outside the clipping bounds and for *δ →* 0, one Newton step gives

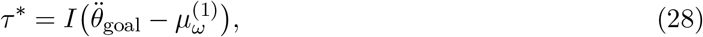

in which the sensory precision cancels between the action gradient and its Hessian. With the regularization constant, the increment is scaled by 1*/*(1 + *δI*^2^*/λ*_vest_): the cancellation is exact for *δ* = 0 and approximate otherwise, and in general a change of *λ*_vest_ alters the increment through this factor. For the parameters used here the ratio *δI*^2^*/λ*_vest_ is 6.125 *×* 10^−4^ at the baseline, 2.45 *×* 10^−3^ when all precisions are scaled by 0.25, and 4.9 *×* 10^−3^ in the global-low regime (1, 1, 1). The increment factor therefore differs from unity by less than 0.5% over the 115 trial settings; the within-trial switches of Section 5.9 leave *λ*_vest_ unchanged, so for them the factor is identical before and after the switch. To this approximation, precision does not act through the action pathway, and belief updating is the route by which precision changes affect postural behavior. At the late steady state, linearizing the body dynamics around upright yields

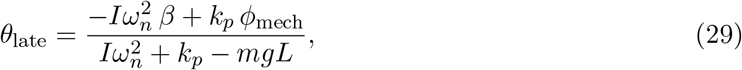

where *θ*_late_ denotes the posture in the late response window. For visual perturbations and sensory-only support tilts (*ϕ*_mech_ = 0) this reduces to

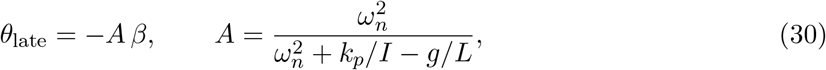

with *A* = 16*/*(16 − 9.81) = 2.585 at *k*_*p*_ = 0 for the parameters in Table 1, decreasing to 1.07 at *k*_*p*_ = 0.9 *mgL*. The denominator of *A* is the restoring stiffness of the linearized closed-loop dynamics (Appendix A.3), yielding the linear stability requirement 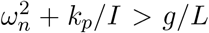: the restorative goal, aided by any intrinsic stiffness, must outpace the gravitational instability. A static sensory goal (*ω*_*n*_ = 0) provides no restorative reference; for *k*_*p*_ *< mgL*, including the *k*_*p*_ = 0 baseline, it violates this requirement and is predicted to fail to maintain stance, whereas an intrinsic stiffness with *k*_*p*_*/I > g/L* would satisfy it without the goal. Equation (30) also fixes the magnitude of the late posture response per unit perturbation at *Aλ*_*t*_*/*Σ*λ* for perturbed-channel precision *λ*_*t*_, giving 0.86 in the equal-precision regime and 0.15 after target-precision reduction at *k*_*p*_ = 0. The nonlinear equilibrium, which replaces *mgLθ* by *mgL* sin *θ*, differs from Eq. (29) by a cubic correction whose coefficient is 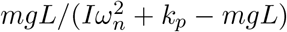 (Appendix A.3); in the trials reported here the difference is below 0.01° for |*θ*_late_| *<* 4° and reaches 0.05° at 8.6° for the baseline goal and 0.12° at 8.3° for *ω*_*n*_ = 3.5.

### 4.4 Reduction ratios: a precision-ratio result and its stiffness limit

For visual perturbations and sensory-only support tilts the amplification factor *A* is common to all precision conditions, so condition-wise posture-shift reductions depend only on precision ratios (Appendix A.4). If the perturbed-channel precision changes from *λ*_*t*_ to 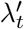, the remaining response fraction is

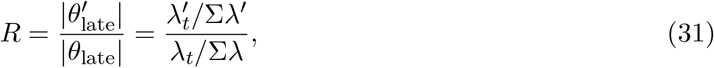

where 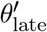 is the late posture under the changed precision; the predicted posture-shift reduction is therefore 1 − *R*, and it is independent of *k*_*p*_. For the single-channel manipulations, *R* = (1*/*17)*/*(8*/*24) = 3*/*17, a predicted reduction of 82.35%; for the equal-amplitude combined conflict with both non-vestibular precisions reduced, *R* = (2*/*10)*/*(16*/*24) = 0.30, a predicted reduction of 70.00%. The same ratio applies to the state-update contributions, so behavioral and mechanism-level reductions are predicted to coincide. By the ratio dependence of Eq. (26), the global low-precision regime is predicted to reduce absolute contributions without reducing the perturbation-driven posture bias.

For a physical support tilt (*ϕ*_mech_ = *ϕ*_sens_ = *ϕ*) the support-tilt term *k*_*p*_*ϕ* in Eq. (29) does not depend on precision, and the remaining fraction becomes

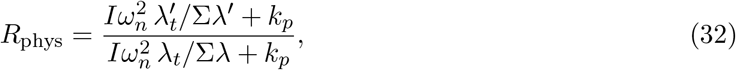

which increases with *k*_*p*_: intrinsic stiffness sets an upper bound on how much sensory reweighting can reduce the response to a physical surface tilt. For the 8-to-1 manipulation, *R*_phys_ rises from 3*/*17 = 0.18 at *k*_*p*_ = 0 to 0.69 at *k*_*p*_ = 0.9 *mgL*, that is, the predicted reduction falls from 82% to 31% for this model setting.

All of the above are linearized, non-saturated, steady-state predictions. Whether the full nonlinear closed loop stands stably, how close its equilibria are to these expressions, how it behaves away from steady state, and whether the pattern survives stochastic observation noise (Appendix A.7) are questions for the simulation experiments that follow.

## 5 Simulation experiments

We tested the predictions of Section 4 in a set of 115 deterministic closed-loop trials using the same body-model equations, action optimizer, and restorative sensory goal. Parameters were fixed within each trial except for the imposed precision schedule of the two precision-switch trials. Unless stated otherwise, sensory perturbations were applied from 2 s to 16 s of an 18 s trial and their effects were evaluated over the late response window 14 s to 16 s (Section 3.8). The trials were organized in blocks that each vary one factor at discrete values. (i) Perturbation-amplitude tests of the predicted gains and curve flattening of Eq. (30): visual offsets and support-surface tilts of ±2, ±4, and ±8 degrees plus the unperturbed case, under equal precision and under target-precision reduction. (ii) Target-precision tests of Eqs. (27) and (30): the target-channel precision *λ*_*t*_ was imposed at 0.5, 1, 2, 2.75, 4, 4.5, 6.25, 8, and 16 with the non-target precisions at 8. (iii) Precision-ratio tests of Eqs. (26) and (31): all precisions scaled together by 0.25 or 4, the global low-precision regime (1, 1, 1), and a combined conflict with *o*_vis_ = +5° and *ϕ* = −5° under (8, 8, 8), (8, 1, 1), and (1, 1, 1). (iv) Intrinsic-stiffness tests of Eqs. (29)–(32): *k*_*p*_ at 0, 0.5, 0.9, and 1.0 *mgL* for visual, sensory-only, and physical support-tilt perturbations, with passive damping *b* at 0, 2, and 20 N m s rad^−1^. (v) Goal-dynamics tests across the stability boundary: *ω*_*n*_ *∈ {*3.0, 3.5, 4.0, 5.0*}* and *ζ∈ {*0.5, 1.0, 1.5*}*, including a small-amplitude (+0.25°) condition near the boundary. (vi) Body-scale tests: *m ∈ {*50, 70, 90*}* kg and *L ∈ {*0.8, 1.0, 1.2*}* m. (vii) Numerical-convergence tests: 16, 32, and 64 D-step iterations with integration steps of 5 and 2.5 ms. Finally, (viii) a within-trial precision switch recorded the time course of the main input and output variables of the D-step and A-step. Blocks (ii)–(vii) used a representative perturbation amplitude of +5°. The complete trial list, with the settling and failure flags defined below, is given in Appendix A.5. Before these blocks, two preliminary checks verified the standing baseline implied by the stability requirement and the signed entry of each perturbation into its intended sensory channel.

The behavioral readout is late posture, *θ*_late_, defined as the mean body tilt *θ* over the late response window. Perturbation gain is the signed response ratio

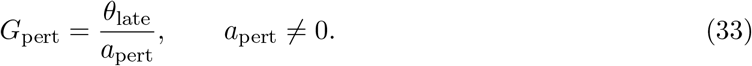

Here, *a*_pert_ is the signed amplitude of the imposed visual or support-surface perturbation. Each simulation was considered stable when the D-step free energy *F*_*D*_ remained finite at every integration step, belief and action variables contained no non-finite values (infinity or NaN), the body model did not diverge (tilt never exceeded 28.6°), and actuator saturation did not dominate the response (torque at its bound on fewer than 5% of integration steps). A trial was considered settled when the mean posture over the late window differed from that over the preceding 2 s by at most 0.01° and the angular velocity in the window had an RMS of at most 0.01 deg*/*s and a maximum of at most 0.05 deg*/*s; only settled trials were compared with the steady-state theory, and agreement was defined as an absolute difference of at most 0.01° from the nonlinear equilibrium (Appendix A.3). Trials that were stable but not settled, diverged, saturated, or had |*θ*_late_| *>* 5° (outside the small-angle regime) are flagged as such and reported. The preliminary checks assess fixed-precision standing and the signed entry of each perturbation into its intended sensory channel.

### 5.1 A stable active inference standing baseline required a restorative generalized goal

The preliminary checks in this and the next subsection used a perturbation interval of 2 s to 6 s with *k*_*p*_ = 0, rather than the longer holds used for steady-state verification. The D-step and A-step remained finite, but a static sensory goal with zero preferred acceleration did not stabilize the closed-loop body model. The reason was structural: action entered the generative model via vestibular angular acceleration, so a static zero-goal could cancel acceleration without creating a restoring-position servo. With this static goal, posture remained unstable despite the physically scaled action mapping of Eq. (16) and the angular-acceleration observation in the vestibular channel. Stable fixed-precision standing was obtained when the preferred generalized observation encoded the return-to-upright acceleration 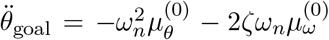. With *ω* = 4.0 and *ζ*= 1.0, the model maintained standing without torque saturation and converged from the initial tilt toward upright. This outcome matches the stability requirement 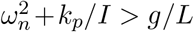 of Section 4: with *k*_*p*_ = 0 the static goal provides no restorative reference and cannot satisfy it, whereas the restorative goal with *ω*_*n*_ = 4.0 does. Section 5.8 tests this requirement directly by crossing the boundary with *ω*_*n*_ = 3.0. This baseline enabled sensory perturbations to be studied without altering the action optimizer or body dynamics.

### 5.2 Controlled sensory probes entered the intended channels

Before testing reweighting, we verified that sensory perturbation conflicts produced the expected signed sensory prediction errors. A positive visual offset increased the visual prediction error, whereas a positive support-surface tilt decreased the proprioceptive prediction error because *y*_prop_ = *θ*−*ϕ*_sens_. In the fixed-precision preliminary probe set, the visual error increased from approximately 0.0125 to 0.089 rad under a positive visual offset, and the proprioceptive error changed from approximately 0.0125 to − 0.078 rad under a positive support-surface tilt. This scale was consistent with the imposed +5° perturbation amplitude (0.087 rad) and with the opposite sign of support-surface tilt in *y*_prop_ = *θ* − *ϕ*_sens_. Here, *ϕ*_sens_ = *ϕ* in the sensory-only condition. These signed-error checks used a probe configuration in which the restorative goal of Eq. (14) was evaluated from the true body state (*θ, ω*) instead of the beliefs. This allowed the entry of each perturbation into its sensory channel to be examined without changing the goal through the posture belief. All reweighting analyses below use the belief-driven goal of Eq. (14).

### 5.3 Context-dependent precision reduced perturbation-driven postural bias

When sensory precision was equal, imposed perturbations strongly biased posture. Under a +5° visual offset, the belief-driven fixed-precision model shifted the late posture by −4.30°. Reducing visual precision from 8 to 1 reduced this shift to −0.76°, an 82.3% reduction. Under a +5° sensory-only support tilt, the equal-precision model shifted posture by +4.30°, and proprioceptive precision reduction reduced the shift to +0.76°, again an 82.3% reduction. In a combined visual-proprioceptive conflict condition (*o*_vis_ = +5°, *ϕ* = −5°), both non-vestibular channels were shifted away from the vestibular body-state signal; reducing both non-vestibular precisions, from equal precision (8, 8, 8) to (8, 1, 1), reduced the posture shift from −8.57° to −2.58°, a 69.8% reduction. Thus, changing only the precision of the unreliable sensory channel was sufficient to change closed-loop postural behavior. These values correspond to the linearized predictions of Eq. (31), 82.35% for a single unreliable perturbation channel and 70.00% for the combined conflict, to within the small-angle approximation (the combined-conflict equilibrium lies at 8.6°, where the sin *θ* nonlin-earity is no longer negligible; the nonlinear equilibria of Appendix A.3 match all four values to within 0.001°). Scaling all precisions together by 0.25 or 4 left every late posture unchanged to four decimal places, as the ratio dependence of Eq. (26) predicts.

### 5.4 State-update gradients identified the mechanism of reweighting

The state-update gradient contribution confirmed that behavioral reweighting originated in belief updating. Visual precision reduction reduced the visual state-update contribution, *C*_vis_, from 0.465 to 0.082 (82.3%), matching the 82.3% reduction in visual perturbation-driven posture shift. Proprioceptive precision reduction produced the same pattern: the proprioceptive state-update contribution, *C*_prop_, decreased by 82.3%, matching the 82.3% behavioral reduction. In the combined visual-proprioceptive conflict condition, the perturbation-channel contribution decreased by 70.0%, matching the 69.8% posture-shift reduction. In every settled trial, *C*_vis_ and *C*_prop_ agreed with the closed form of Eq. (27) to within 0.005%. Two simpler summaries were considered but not used as primary indices. Normalized contribution shares, which divide each channel’s contribution by the summed contribution across channels, can remain approximately conserved when the belief redistributes prediction errors across channels. *λ*|*ε*| scores multiply scalar precision by absolute prediction error but do not account for the observation-model Jacobian. The closed-form expression of Eq. (27) reproduced the target-precision gradient contributions of Section 5.6 to four decimal places.

### 5.5 Precision control reproduced Peterka-style perturbation-response curves

The perturbation-amplitude tests showed the qualitative hallmark of sensory reweighting: target-channel precision reduction flattened the perturbation-response curve without changing the body dynamics, action optimizer, or restorative goal (Fig. 3; Table 2). For visual offsets, equal precision produced a perturbation gain of −0.86 at ±2° and ±4° (−0.858 at ±8°), whereas visual precision reduction reduced the gain magnitude to 0.152. For support tilt, equal precision produced a perturbation gain of +0.86, whereas proprioceptive precision reduction reduced the gain to +0.152. These gains match the predicted magnitudes *Aλ*_*t*_*/*Σ*λ* = 0.861 and 0.152 of Eq. (30); the only departures from linearity are the sin *θ* correction at ±8° (0.1°), which the nonlinear equilibrium reproduces to 0.0001°. Evaluating the same trials over the earlier 5 s to 6 s window gives gains about 5% smaller because of a residual closed-loop transient with a time constant of about 1 s (Appendix A.5). Across the nonzero perturbation amplitudes, *C*_vis_ decreased by 82.35% in the visual-offset tests and *C*_prop_ decreased by 82.35% in the support-surface-tilt tests.

**Table 2:** Perturbation gain and state-update contribution for the perturbation-amplitude tests (±2, ±4, ±8°; linear predictions in parentheses).

| Perturbation | Equal precision | Target precision 8 → 1 | *C*<sub>target</sub> reduction |
|---|---|---|---|
| Visual offset | \|*G*<sub>pert</sub>\| = 0.858–0.861 (0.861) | \|*G*<sub>pert</sub>\| = 0.152 (0.152) | 82.35% (82.35%) |
| Support-surface tilt | \|*G*<sub>pert</sub>\| = 0.858–0.861 (0.861) | \|*G*<sub>pert</sub>\| = 0.152 (0.152) | 82.35% (82.35%) |

**Figure 3.**
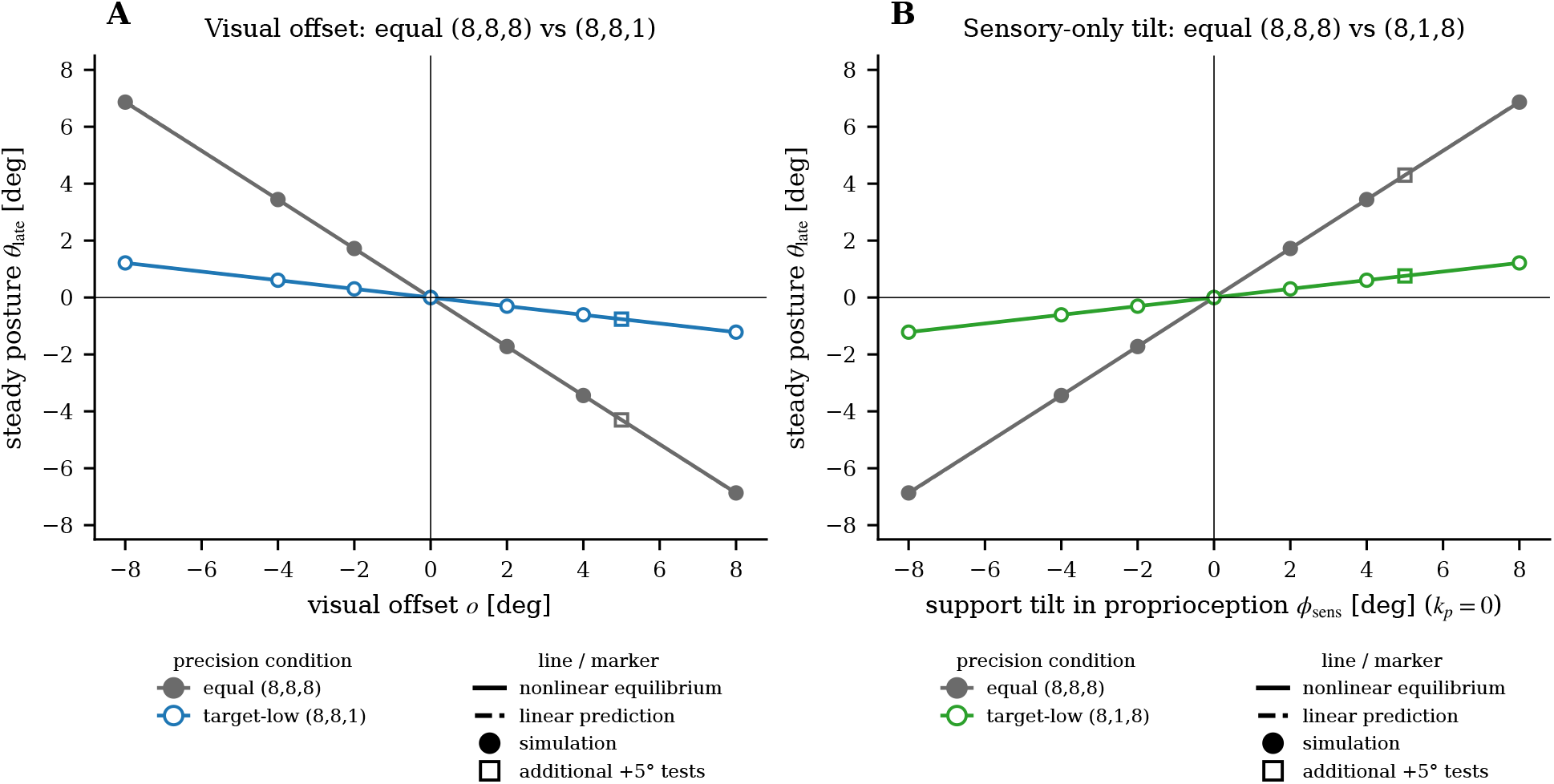
Precision control reproduces Peterka-style flattening of perturbation-response curves. (A) Visual offset. (B) Sensory-only support-surface tilt (*k*_*p*_ = 0). Each marker is a separate trial with fixed precision (equal, (8, 8, 8); or target-low, (8, 8, 1) in A and (8, 1, 8) in B), perturbation applied from 2 to 16 s and posture averaged over 14–16 s; open squares are additional +5° trials under the same protocol. Solid lines are the nonlinear equilibrium (Appendix A.3) and dashed lines the linear prediction of Eq. (30); the two nearly overlap at this scale (they differ by less than 0.3° at 8°) but are not identical. Lines are theory, markers are simulations; deterministic trials carry no error bars.

### 5.6 Imposed target-channel precision controlled contribution and posture as predicted

Imposing the target-channel precision directly at nine values between 0.5 and 16 produced monotonic control of state-update contribution and posture (Fig. 4). For a visual offset, the visual contribution increased from 0.042 at *λ*_*t*_ = 0.5 to 0.698 at *λ*_*t*_ = 16, passing through 0.082, 0.155, 0.205, 0.279, 0.307, 0.392, and 0.465 at *λ*_*t*_ = 1, 2, 2.75, 4, 4.5, 6.25, and 8; every value matched the closed form of Eq. (27) to four decimal places (the sublinear growth reflects the factor (Σ*λ*−*λ*_*t*_)*/*Σ*λ*). The proprioceptive tests showed the same pattern with the opposite sign of posture. Late posture followed the nonlinear equilibrium to within 0.0001° at all nine values, and the linear prediction to within 0.02° except at *λ*_*t*_ = 16, where the 6.4° equilibrium lies outside the small-angle regime. All of these trials were stable and settled. Precision was imposed directly as the condition parameter, without assuming a functional mapping from environmental reliability to precision.

**Figure 4.**
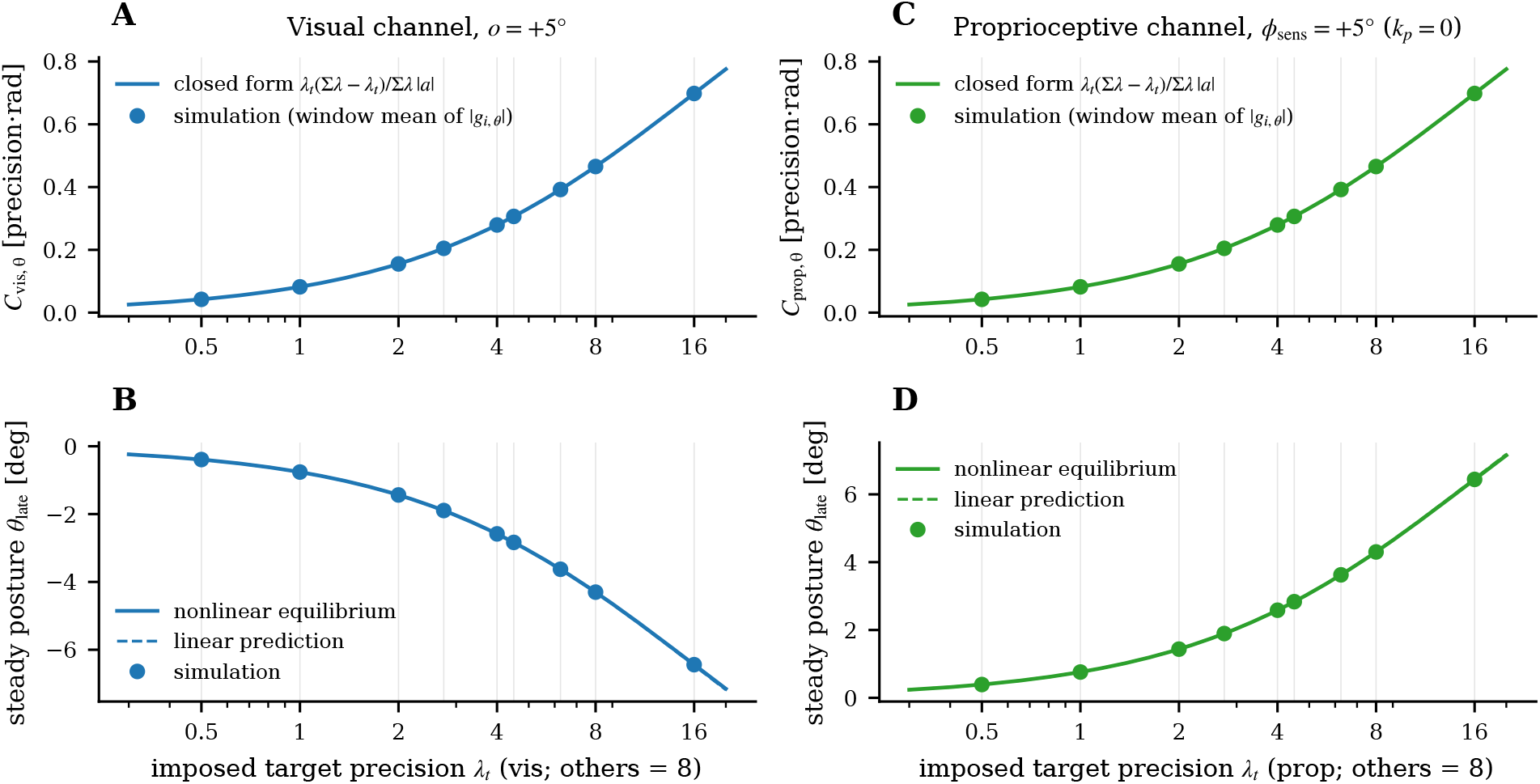
Imposed target-channel precision controls contribution and posture as predicted in closed form. (A) Visual state-update contribution *C*_vis_ and (B) late posture for a +5° visual offset as the visual precision *λ*_*t*_ is imposed at nine values from 0.5 to 16 (log axis; non-target precisions 8). (C, D) The same for the proprioceptive channel under a +5° sensory-only support tilt. Lines: closed form of Eq. (27) (A, C), nonlinear equilibrium (solid) and linear prediction (dashed) of Eq. (30) (B, D). Markers: simulations.

### 5.7 Control analyses defined the scope of the prediction

The control analysis distinguished channel-selective sensory reweighting from global precision reduction (Fig. 5). For a +5° visual offset, target-selective visual precision reduction produced a small posture shift (−0.76°) and a perturbation-channel contribution of 0.082. Equal precision produced a larger shift (−4.30°) and a contribution of 0.465. Global low precision (1, 1, 1) reduced the absolute contribution to 0.058, but posture still shifted by −4.30°, the same behavioral bias as equal precision to four decimal places. The proprioceptive condition showed the same structure: target-selective proprioceptive precision reduction produced a small shift (+0.76°), whereas equal precision and global low precision produced the same larger shift (+4.30°). Therefore, absolute contribution magnitude alone is insufficient to identify reweighting. Reweighting requires a context-selective change in relative precision, and a behavioral posture shift is necessary to distinguish it from global uncertainty. The ratio dependence of the bias in Eq. (26) explains this result: global low precision preserves the same precision ratios as equal precision, so it reduces absolute contribution magnitude without removing the perturbation-induced belief bias, exactly as predicted in Section 4. The model prediction is therefore specific: lowering all sensory precisions by a common factor leaves the perturbation-driven posture bias unchanged while reducing every channel’s update pressure in proportion.

**Figure 5.**
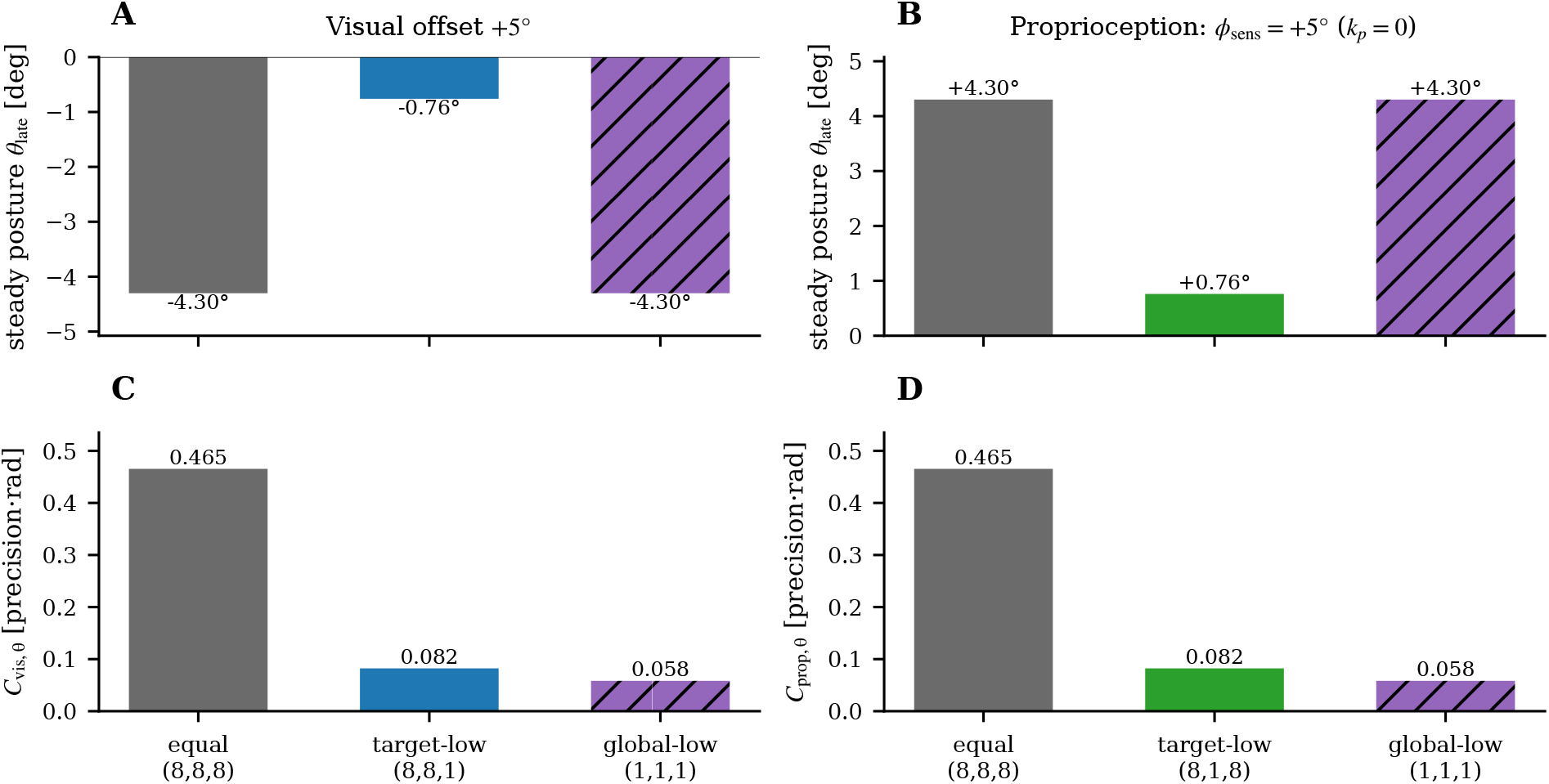
Global precision reduction does not reweight. (A, B) Late posture and (C, D) target-channel contribution for a +5° visual offset (A, C) and a +5° sensory-only support tilt (B, D) under equal precision, target-precision reduction, and global low precision (1, 1, 1) (hatched), all under the same long-hold protocol. Target-precision reduction reduces both posture shift and contribution; global low precision reduces the contribution to 0.058 but leaves the posture shift unchanged, because precision ratios, and hence the belief bias of Eq. (26), are unchanged. A small contribution is therefore not sufficient for a small behavioral bias. Deterministic trials; no error bars.

### 5.8 Intrinsic ankle stiffness changed the amplification but not the precisionratio reduction, except for physical surface tilts

Fig. 6 tests the predictions of Eqs. (29)–(32) by adding intrinsic ankle stiffness at 0.5, 0.9, and 1.0 *mgL*. For a +5° visual offset (Fig. 6A), the equal-precision posture shift decreased from −4.30° at *k*_*p*_ = 0 to −2.40°, −1.78°, and −1.67°, following the reduced amplification factor *A*, and the target-low shift decreased from −0.76° to −0.42°, −0.31°, and −0.29°; the remaining fraction *R* was 0.1767 at *k*_*p*_ = 0 and 0.1765 at the three nonzero stiffnesses (Fig. 6C), against the linearized value 3*/*17 = 0.1765 of Eq. (31) at every *k*_*p*_; the small departure at *k*_*p*_ = 0 is the sin *θ* correction at the larger equilibrium there. Sensory-only support tilts gave the same values with opposite sign. For a physical support tilt (Fig. 6B), the support-tilt torque *k*_*p*_*ϕ* pulled the body with the surface irrespective of precision: the equal-precision shift grew slightly (+4.61°, +4.71°, +4.73°), the target-low shift grew substantially (+2.63°, +3.25°, +3.36°), and the remaining fraction rose from 0.18 to 0.57, 0.69, and 0.71, matching *R*_phys_ of Eq. (32) (0.571, 0.690, 0.710). At *k*_*p*_ = 0 the physical and sensory-only tilts coincided exactly, confirming that the baseline support-tilt condition is purely sensory. The passive damping *b* had no effect on any late posture (*b* = 0, 2, and 20 N m s rad^−1^ gave identical values to four decimal places at *k*_*p*_ = 0 and 0.9 *mgL*), consistent with its absence from the equilibrium equations; it does contribute to the transient (Appendix A.3). Finally, intrinsic stiffness restored stability across the goal-dynamics boundary (Fig. 6D): with *ω*_*n*_ = 3.0, for which 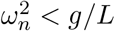, the *k*_*p*_ = 0 model diverged within 12 s with the torque at its bound, whereas *k*_*p*_ = 0.9 *mgL* satisfied 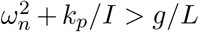 and settled at −1.87°, as predicted. Thus, omitting intrinsic stiffness changes the absolute size of the perturbation responses but leaves the precision-ratio reduction for visual and sensory-only perturbations unchanged in the linearized theory and approximately unchanged in the simulations; for physical surface tilts, intrinsic stiffness bounds the reduction that reweighting can achieve. The quantitative values (82% *→* 31%) hold for this model setting (*ω*_*n*_ = 4, *ζ*= 1, +5°, precision 8 to 1) and are not proposed as universal proportions.

**Figure 6.**
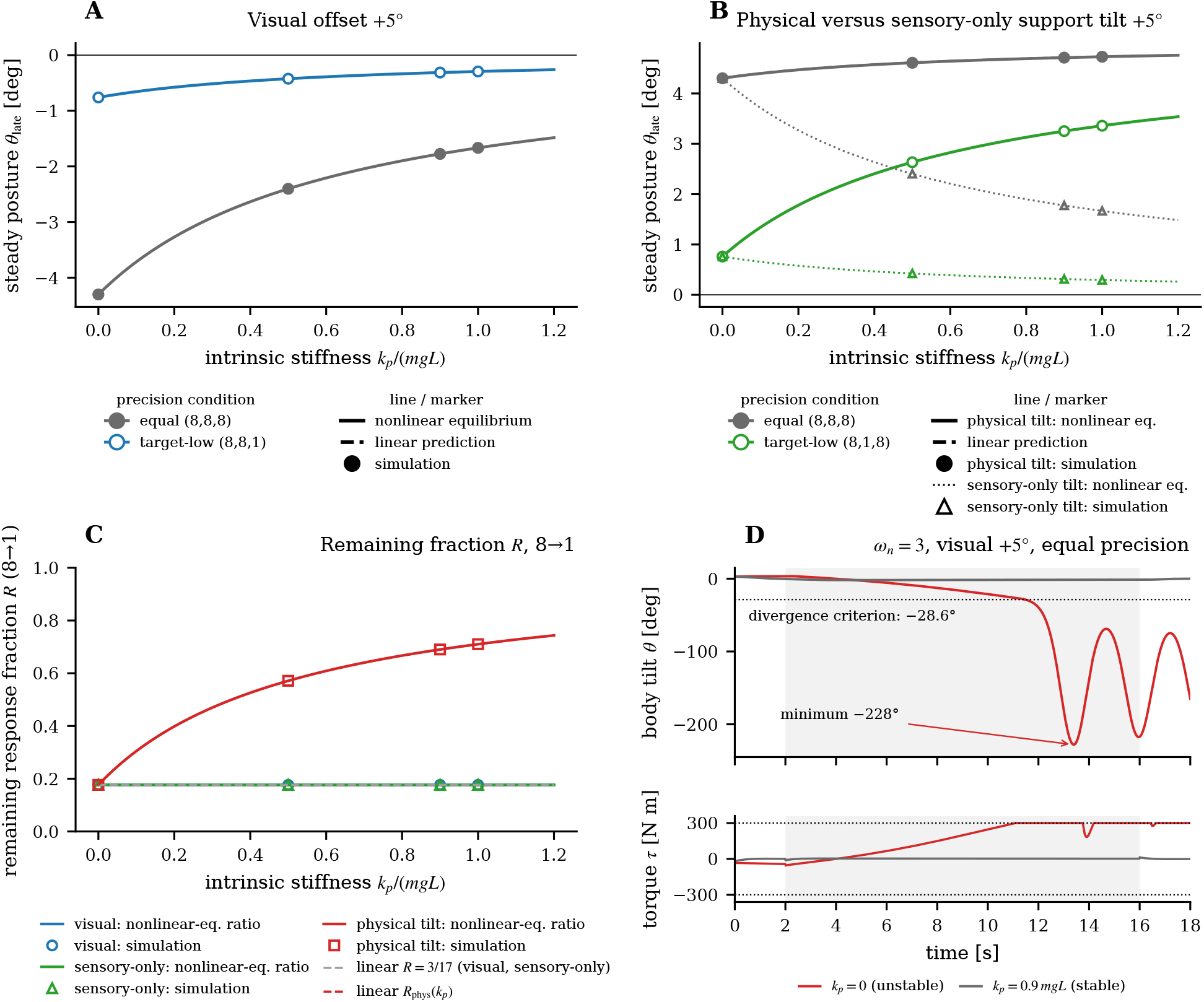
Intrinsic ankle stiffness. (A) Late posture for a +5° visual offset versus *k*_*p*_*/*(*mgL*) under equal (grey) and target-low (blue) precision. (B) Late posture for a +5° physical support tilt (*ϕ*_mech_ = *ϕ*_sens_; solid lines, circles) and for the sensory-only tilt (*ϕ*_mech_ = 0; dotted lines, triangles) under equal (grey) and target-low (green) precision. (C) Remaining response fraction *R* for the 8-to-1 precision reduction. Dashed: linearized predictions *R* = 3*/*17 (visual and sensory-only) and *R*_phys_(*k*_*p*_) of Eq. (32); solid: ratio of the nonlinear equilibria; markers: simulations. The visual and sensory-only series coincide. (D) Body tilt for *ω*_*n*_ = 3.0 with *k*_*p*_ = 0 (red; predicted unstable) and *k*_*p*_ = 0.9 *mgL* (grey; predicted stable), equal precision, +5° visual offset from 2 to 16 s (shaded). Upper: complete body-tilt traces; lower: control torque. Dotted lines mark the −28.6° divergence criterion and the ±300 N m torque bounds. The *k*_*p*_ = 0 trial crosses the divergence criterion and reaches a minimum of −228°; its subsequent numerical trajectory is outside the quiet-standing regime.

### 5.9 A within-trial precision switch showed the belief-to-action time course

Fig. 7 shows the main model variables when the target-channel precision is switched within a trial while the belief, torque, and body state are carried across the switch: the perturbation is applied at 2 s, the target precision is reduced from 8 to 1 at 7 s and restored at 12 s, and the perturbation is removed at 17 s. At a fixed belief and observation, switching the target precision from 8 to 1 scales the target-channel gradient by one eighth and leaves the other channel gradients unchanged; this was verified by evaluating the gradient function under both precisions at the switch step. In the recorded time course, the D-step also changes the belief during the switch step, so the prediction errors and hence all channel gradients change at that step: between the samples at 6.995 s and 7.000 s the target gradient fell to 0.147 of its previous value and the vestibular and non-target gradients to 0.652 (Fig. 7C). The tilt belief then moved toward the vestibular and non-target evidence over several numerical update steps (Appendix A.6), the preferred acceleration of Eq. (14) changed accordingly (Fig. 7D), and the torque followed the goal (Fig. 7E); the body then relaxed from −4.26° toward the constant-low-precision level of −0.76° with a 10–90% response time of 2.36 s (Table A.2). Restoring the precision and removing the perturbation produced transitions of opposite direction with 10–90% times of 2.37 s and 2.38 s. The proprioceptive switch (Fig. 7, right) behaved identically with the opposite sign of posture.

**Figure 7.**
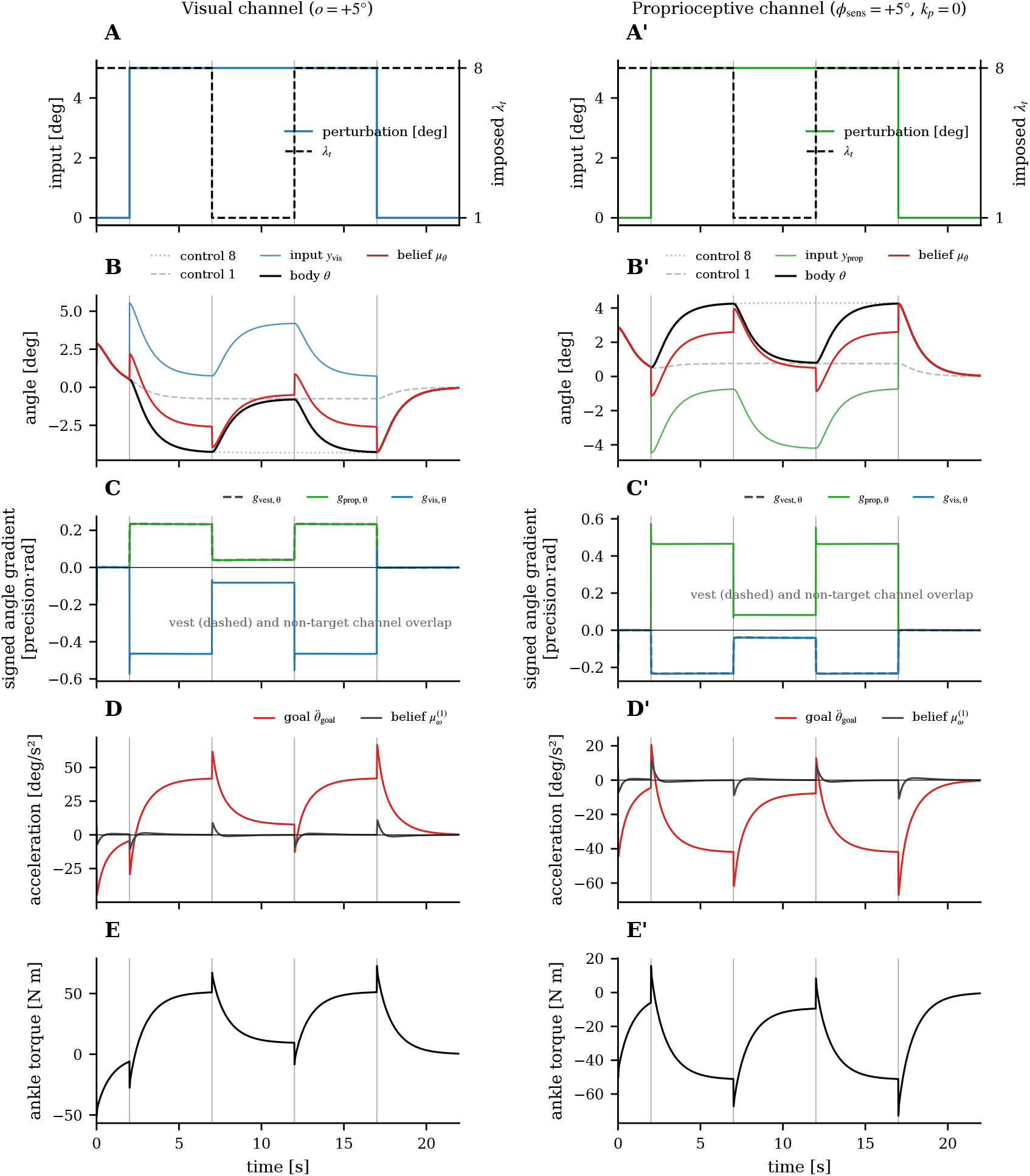
Within-trial precision switch. Left column: +5° visual offset with the visual precision switched; right column: +5° sensory-only support tilt (*k*_*p*_ = 0) with the proprioceptive precision switched. Vertical lines mark perturbation onset (2 s), the 8-to-1 switch (7 s), the 1-to-8 switch (12 s), and perturbation offset (17 s). (A) Perturbation input and the imposed target precision *λ*_*t*_. (B) Perturbed-channel input, body tilt, and tilt belief 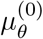 after the D-step; grey lines are control trials with constant precision 8 (dotted) and 1 (dashed). (C) Signed angle gradients *g*_*i,θ*_ of the three channels; the belief moves along −*g*; the vestibular (dashed) and non-target channel overlap. (D) Preferred acceleration 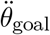 and the belief 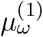. (E) Ankle torque. Within each 5 ms step the record order is observation, D-step (belief shown), A-step (torque shown, applied to the following integration), and body integration; the body state shown precedes integration.

Two features of this time course bear on the mechanism. First, at the switch step the A-step objective also changes, but the Newton increment computed from the same beliefs under the old and the new precision was identical: because only a non-vestibular precision was switched and the vestibular precision *λ*_vest_ that scales the action-dependent term stayed at 8, the direct action pathway contributed nothing to the change in torque, which arose entirely through the belief (Section 4). This observation is specific to switches that leave *λ*_vest_ unchanged; with *δ >* 0 a change of *λ*_vest_ would alter the regularization factor. Second, the non-zero steady goal acceleration during the perturbation (Fig. 7D) is what produces the steady torque that balances gravity at the biased posture; it is the sensory-level expression of the set point discussed in Section 3.5. The response times are indices of the model’s closed-loop dynamics under an imposed precision schedule; they are not estimates of human reweighting time constants or neural delays.

### 5.10 Range of verification

Of the 115 trials, 111 settled, and every settled trial agreed with the nonlinear equilibrium of Appendix A.3 to within 0.0006° (Appendix A.5, Fig. A.1). Twelve trials carried the large-angle flag |*θ*_late_| *>* 5°: ten settled trials (the ±8° amplitudes, *λ*_*t*_ = 16, the two combined-conflict trials with equal and global-low precision, *ω*_*n*_ = 3.5 at +5°, and *L* = 0.8 m), for which the linear prediction deviates by up to 0.12° while the nonlinear equilibrium still holds, and the two unsettled *ω*_*n*_ = 3.0 trials described next. The four trials that did not settle were the two *ω*_*n*_ = 3.0, *k*_*p*_ = 0 trials, for which the loop is predicted unstable (one diverged with saturated torque; the other, with the target precision reduced, had a mean tilt of 5.5° in the 14–16 s evaluation window and reached 7.4° at 18 s without crossing the divergence threshold within the observation interval), and the two precision-switch trials, whose evaluation window follows the restoring switch by only 5 s and which are reported as time courses rather than steady states. Body mass had no effect on late posture (*m* = 50, 70, 90 kg gave identical values, as *τ*^*\**^*/I* cancels *m*), whereas center-of-mass height changed the amplification through *g/L* (−7.08° at *L* = 0.8 m and −3.41° at *L* = 1.2 m for the equal-precision visual condition), and varying the goal damping ratio *ζ*changed the late posture by less than 0.001°. One near-boundary condition (*ω*_*n*_ = 3.2, linearized slow time constant 14.7 s) was excluded from the main run by the pre-specified 60 s hold cap because 10 *τ*_slow_ exceeded it; a pilot run of that condition with a 98 s hold differed from the linear prediction by 0.014° and from the nonlinear equilibrium by 0.005° in its final window, and its steady state is not claimed as verified. The 115 trials are sparse verification points, not a map of the stable region; the analytical boundary is Eq. (30).

## 6 Discussion

The present results support a minimal mathematical account of sensory reweighting in quiet standing. In the model, context did not select a new controller or retune the feedback gain. Instead, context changed the relative precision of sensory prediction errors. This precision change altered how strongly each perturbation channel updated the posture belief, and the altered belief propa-gated through the same free-energy-minimizing action loop to produce different postural responses. The action update did not provide a separate precision-dependent pathway for reweighting: as shown in Section 4 (Eq. (28)), the sensory precision cancels from the Newton action update exactly for *δ* = 0 and, for the regularized update used here, up to a factor 1*/*(1 + *δI*^2^*/λ*_vest_) that differs from unity by less than 0.5% over the tested settings, so belief updating is the route through which precision changes produce reweighting in this model. The within-trial switch (Section 5.9) showed this directly for switches of the visual or proprioceptive precision: the action increment computed from the same beliefs was identical before and after the switch, and the torque changed only as the belief changed.

The intrinsic-stiffness analysis distinguishes the effects of intrinsic ankle stiffness on absolute responses and on reweighting. Intrinsic stiffness adds to the closed-loop restoring stiffness, reduces the amplification of belief bias into posture, and can stabilize a goal that would otherwise be too weak (Section 5.8). For visual perturbations and for support tilts that act only through proprioception, the precision-ratio reduction is unchanged in the linearized theory, and approximately unchanged in the simulations, because the stiffness multiplies both conditions alike. For physical support tilts, however, the spring torque pulls the body with the surface irrespective of what the nervous system believes, so reweighting can reduce only the actively generated part of the response; Eq. (32) gives this bound in closed form. This distinction between the support-tilt reference in proprioception and the physical support tilt is a prediction that posturography with and without mechanical coupling could test. Passive damping, by contrast, affects only the transient and is dominated by the goal’s damping term. In the delayed feedback model of Le Mouël and Brette (2019), increases in intrinsic stiffness require coadaptation of sensorimotor gain to maintain stability; the present model has no delay, so it exhibits only the stabilizing side of that trade-off, and intrinsic stiffness in standing humans is itself variable with sway, torque, and individual (Sakanaka et al., 2018, 2021), which the fixed *k*_*p*_ used here does not represent.

The model’s goal law corresponds to a stiffness factor of 1120 N m rad^−1^ and a damping factor of 560 N m s rad^−1^ acting on the believed tilt and velocity, plus an acceleration-belief term (Section 3.5); the net restoring stiffness after subtracting the gravitational load is 433 N m rad^−1^ at *k*_*p*_ = 0. These values were chosen to give a stable baseline, not fitted to human data. Within each intrinsic-stiffness comparison, the goal gains were held fixed as *k*_*p*_ changed. This isolates the mechanical effects on stability and posture; it does not quantify reductions in human muscle activation, neural drive, or metabolic cost. A reduction of active inference to proportional-integral-derivative control, with precision parameters related to controller gains, has been derived under specific linear-model and approximation assumptions (Baltieri and Buckley, 2019); this does not establish that every linear active-inference implementation has that form. The mathematical correspondence to an observer-plus-feedback controller is intended to make the model comparable, not to claim that active inference controls stance better than, or through a different physical pathway from, such a controller, nor to exclude intermittent or predictive control schemes (Loram et al., 2022). The specific content of the present model is that the sensory weights enter one objective that also selects the action.

The gradient-based contribution measure is important for interpreting this result. If reweighting were assessed solely by the experimenter-specified precision values, the result would be circular. If it were assessed only by normalized prediction-error shares, the result could be missed because the belief can redistribute errors across channels. The absolute state-update gradient contribution addresses these two problems by measuring the actual pull of a channel’s prediction error on the posture belief. However, as the global-low-precision control analysis shows, the absolute gradient magnitude is not, by itself, sufficient to identify reweighting. Its close quantitative match to the behavioral posture-shift reduction is what indicates that precision-controlled belief updating is the mechanism through which reweighting affects action in this model.

The global low-precision control condition clarifies the model prediction that lowering all sensory precisions by a common factor reduces every channel’s update pressure but does not change the relative balance among channels. Consequently, the perturbed channel still biases posture by the same amount as in the equal-precision regime. Thus, sensory reweighting should not be identified with low precision per se. It is a relative and context-selective redistribution of precision across sensory channels.

This study has several limitations. The body model is a one-link inverted pendulum, not a full musculoskeletal model. The action is a single ankle torque, not muscle activation. The vestibular, proprioceptive, and visual observation equations are deliberately minimal. The model contains no sensorimotor delay: feedback models of human balance include a physiologically significant effective loop delay (Peterka, 2002; Peterka et al., 2018). Its estimate depends on stimulus and model: Peterka et al. (2018) reported mean delays of 120 ms across three test types involving support-surface tilt and 206 ms for visual-stimulus tests. These are effective model parameters, not direct measurements of a single neural conduction delay. Robot-imposed delays also modify vestibular-evoked responses, with partial recovery after training (Rasman et al., 2021). The absence of delay here means that the stability condition of Section 4, the closed-loop time constants, and the transient responses of Section 5.9 cannot be extrapolated to human stance; introducing delay requires a new stability and transient analysis. The model also does not specify how channel precisions are determined in humans. In biological postural control, such reliability assignments may depend on sensory noise, cue validity, task relevance, attention, learning, and externally applied balance disturbances; here they are imposed to isolate their closed-loop consequences, and the within-trial switch is an imposed schedule, not a model of online reliability estimation. The model therefore does not predict the time course of reweighting measured in humans. Such experiments report transition delays, fast and slow components, and condition-dependent asymmetries, with different dynamics for stimulus-evoked sway and sway variability (Assländer and Peterka, 2014, 2016; Honeine et al., 2015); the response times reported here are properties of the goal dynamics and of the numerical belief-update schedule. Their numerical resemblance to an experimental adjustment time would not establish a shared physiological mechanism. The perturbation-response curves are linear within each precision regime up to the sin *θ* correction; the model does not attempt to reproduce the experimentally measured amplitude-dependent gain reduction reported in human posturography. The Peterka-style comparison in this paper is therefore limited to between-regime flattening induced by changes in precision. The support surface is modelled as a static tilt; surface motion and velocity-dependent passive torques are not represented, so platform-rotation responses at finite frequency are outside the model’s scope. Long holds were used only to verify the steady-state theory; near the stability boundary the loop settles slowly, and one condition whose predicted hold exceeded the cap was excluded, which limits the parameter range over which the equilibrium predictions were verified (Section 5.10). Precision should be read primarily as a dimensionless inverse-uncertainty parameter of the generative model rather than as an empirically estimated noise variance. However, a stochastic robustness check in which channel precisions were paired with corresponding observation-noise variances preserved the same reweighting pattern under the same long-hold protocol (Appendix A.7). The model explains sensory reweighting at the level of a minimal computational mechanism, namely relative precision changes propagating through precision-weighted belief updating to closed-loop postural action, not as an identification of the internal model used by the nervous system. In particular, the absence of an inverted-pendulum dynamics prior in the D-step is a modeling restriction used to isolate the precision mechanism; it is not a claim that human postural control lacks internal body-dynamics models. Human stance control likely combines internal dynamics, reflex pathways, body mechanics, and continuity constraints in a more distributed way. The results should therefore be interpreted as a mechanistic proof of concept of multisensory reweighting in postural control, not as a complete model of human sensorimotor physiology.

Future work should extend the model to multi-joint stance, muscle-level actuation, sensorimotor delays, and richer sensory contexts. It should also compare the model more directly with human posturography data, including fitted reweighting curves, individual differences, and condition-specific estimates of sensory reliability. A further direction is to model how relative precision parameters are inferred or adapted from sensory statistics and task context, and to integrate this precision-control mechanism into locomotor active inference models that account for changes in sensory reliability across contact phases and terrain conditions.

## 7 Concluonsion

A minimal continuous-time mathematical model reproduced key signatures of postural sensory reweighting. Fixed-point analysis yielded closed-form predictions for belief bias, state-update contributions, steady posture, and posture-shift reductions, together with a linear stability condition. These predictions specify how the outcomes depend on channel precisions, goal dynamics, and intrinsic ankle stiffness. Closed-loop simulations matched the equilibrium predictions in all 111 trials that settled. The two conditions predicted to be unstable did not settle. Channel-selective precision control reduced perturbation-driven posture shifts, flattened Peterka-style perturbation-response curves, and controlled sensory contribution monotonically over a 32-fold range of imposed target precision. Intrinsic ankle stiffness changed the size of the responses but left the precisionratio reduction for visual and sensory-only perturbations unchanged to within the linearization, and bounded that reduction for physical support tilts. Control analyses showed that reweighting is not a global reduction in precision, but a context-selective change in relative precision across sensory channels, and a within-trial precision switch showed the change reaching ankle torque through the belief. The model has no sensorimotor delay and does not estimate precision online; within those limits, these findings support relative precision control as a compact mathematical account of adaptive sensory reweighting in quiet standing.

## Statements and Declarations

### Ethics approval

Not applicable. This study is based entirely on numerical simulation and involved no human participants and no animals.

### Consent to participate

Not applicable.

### Consent to publish

Not applicable.

### Data availability

The model implementation, simulation and analysis scripts, trial manifest, per-trial summary results, data underlying the numerical figures, and provenance records are archived on Zenodo at 10.5281/zenodo.23966197 (release v0.3.0-bc-r1). The code repository is available at github.com/jkoba0512/vst-aif-posture. Raw simulation time series are not included in the archive but can be regenerated using the scripts.

### Author contributions

Jun Kobayashi is the sole author and performed the conceptualization, methodology, software, validation, formal analysis, investigation, data curation, and visualization, and wrote and revised the manuscript.

### Preprint

This manuscript is an updated version of the bioRxiv preprint posted at https://doi.org/10.64898/2026.06.23.733972.

### Declaration of generative AI and AI-assisted technologies in the manuscript preparation process

During the preparation of this work, the author used Claude (Anthropic) and Codex (OpenAI) for language editing, manuscript revision, checking of analytical derivations, code-review assistance, and submission workflow support. The author reviewed and edited the output as needed and takes full responsibility for the content of the published article. No AI tool is listed as an author.

## A Derivations and stochastic robustness check

This appendix derives the fixed-point results stated in Section 4 for the linearized constant-perturbation regime used in the late response window, together with the linear stability condition of the closed loop. Let

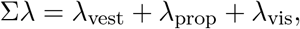

and let *o* denote a constant visual offset and *ϕ* a constant support-surface tilt. With the observation definitions in the Methods, the order-0 angle-channel signals are *θ, θ* − *ϕ*, and *θ* + *o* for vestibular, proprioceptive, and visual channels, respectively.

### A.1 D-step fixed point

The D-step update in Eq. (13) is gradient descent on *F*_*D*_, so a belief is a fixed point of the update exactly when 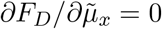, that is, when the partial derivative of *F*_*D*_ with respect to every belief coordinate vanishes. The derivations below compute these stationarity conditions coordinate by coordinate: for each belief coordinate, they collect the terms of *F*_*D*_ that contain that coordinate and set the corresponding partial derivative to zero. Because the observation model *g* in Eq. (11) is linear in the belief and the free-energy objective is quadratic, each stationarity condition is a linear equation with a closed-form solution; since every belief coordinate carries a positive precision weight, the stationary point is the unique minimizer of *F*_*D*_. Each sensory channel is predicted by a single belief coordinate, and the smoothness term contains no order-0 component, so the order-0 tilt belief 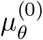 enters *F*_*D*_ only through the three order-0 angle-channel terms,

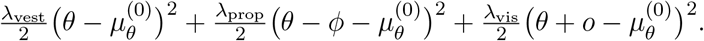

Setting 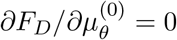 gives

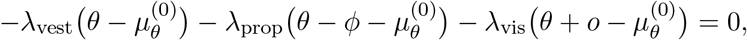

whose solution is the precision-weighted sensory average:

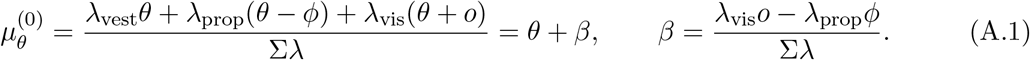

Thus, the perturbation-induced belief bias *β* depends on relative sensory precision. The state smoothness precision 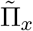 does not enter Eq. (A.1): the smoothness term 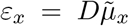 penalizes generalized motion and does not directly constrain the order-0 constant offset.

The remaining fixed points are obtained using the same coordinate-wise procedure. With 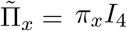, the smoothness term contributes 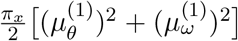to *F*_*D*_. The order-0 angular-velocity belief appears in a single sensory term, 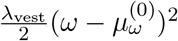, so its stationarity condition directly gives

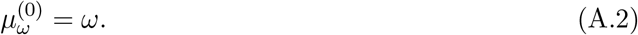

Under constant perturbations, the three first-order angle-channel observations all equal *ω* and are predicted by 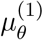, whereas only the vestibular channel observes stationarity conditions are 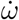, predicted by 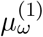; their

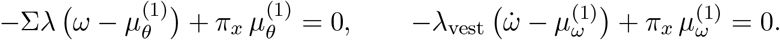

Solving these two linear equations gives

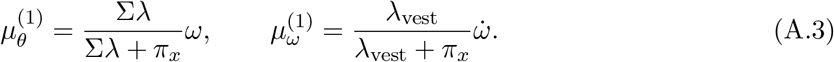

Each first-order fixed point is thus a precision-weighted average of its sensory evidence and the zero prior mean implied by the smoothness term, which acts as a zero-mean Gaussian prior with precision *π*_*x*_ on the first-order components.

### A.2 State-update contribution

At the fixed point, the order-0 channel contribution is the channel precision multiplied by its residual conflict with the belief. For a visual offset with *ϕ* = 0,

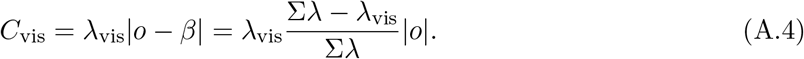

For a support-surface tilt with *o* = 0,

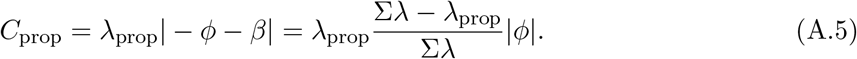

For the visual target-precision tests with *o* = 5° = 0.0873 rad and *λ*_vest_ = *λ*_prop_ = 8, Eq. (A.4) gives the values in Table A.1.

The global low-precision condition (1, 1, 1) gives *C*_vis_ = 0.0582, matching the simulated value of 0.058. Because Eq. (A.1) depends on precision ratios, global low precision preserves the same perturbation-induced belief bias as equal precision while reducing the absolute gradient magnitude. The fixed point also fixes the contribution shares. For a single perturbed channel, the non-perturbed channels share the residual −*β*, so for a visual offset their order-0 contributions sum to

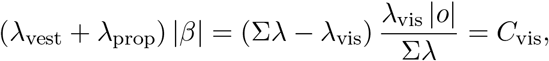

and symmetrically for a support-surface tilt. The perturbed channel’s share of the total order-0 contribution is therefore exactly 1*/*2 at the fixed point, for any precision values. Normalized shares are thus invariant at the fixed point and cannot register reweighting, which motivates the absolute index *C*_*i*_.

### A.3 Closed-loop posture bias

Up to terms that are constant with respect to *τ*, the scalar A-step minimizes 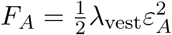 with

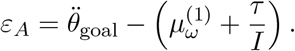

Since

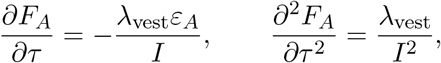

neglecting the small regularization constant *δ*, one Newton step gives

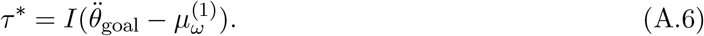

The sensory precision cancels from this expression, so precision affects action through the belief produced by the D-step, not through a separate precision gain in the A-step. With the regularization constant retained, the Newton increment isΔ*τ* = *Iε*_*A*_*/*(1 + *δI*^2^*/λ*_vest_), so the cancellation holds to relative accuracy *δI*^2^*/λ*_vest_ (6 *×* 10^−4^ for *δ* = 10^−6^, *I* = 70, *λ*_vest_ = 8).

At the late steady state, 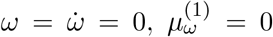, and 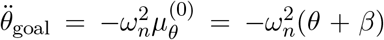. The equilibrium condition of the body dynamics with intrinsic stiffness, Eq. (2), is then

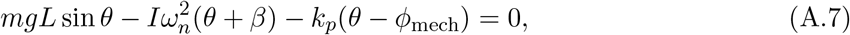

whose root near upright is the nonlinear equilibrium used as the reference for all simulation comparisons. Linearizing sin *θ ≈ θ* Yields

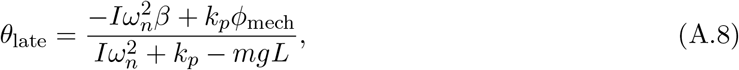

and, for *ϕ*_mech_ = 0,

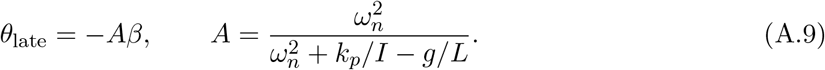

With *ω*_*n*_ = 4.0, *L* = 1.0 m, and *k*_*p*_ = 0, *A* = 16*/*(16 − 9.81) = 2.585; with *k*_*p*_ = 0.9 *mgL, k*_*p*_*/I* = 0.9 *g/L* = 8.83 s^−2^ and *A* = 1.065. Subtracting Eq. (A.8) from Eq. (A.7) gives, with 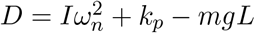,

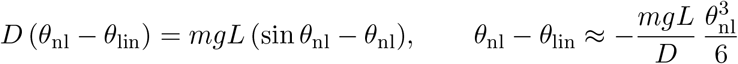

in radians, so the nonlinear equilibrium is slightly closer to upright than the linear one. The coefficient *mgL/D* is 1.58 at the baseline (*ω*_*n*_ = 4, *k*_*p*_ = 0), 4.02 at *ω*_*n*_ = 3.5, and 0.65 at *k*_*p*_ = 0.9 *mgL*. In the simulations the correction is 0.006° at 4.30° (baseline, +5° visual offset), 0.05° at 8.57° (baseline, combined conflict), and 0.11° at 8.25° (*ω*_*n*_ = 3.5, +5° visual offset), each within 0.001° of the cubic estimate.

The stability requirement follows from the closed-loop dynamics rather than from the equilibrium expression alone. Treating the belief as equilibrated on the fast D-step timescale, so that 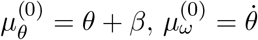, and 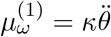 with *κ* = *λ*_vest_/(*λ*_vest_ + *π* _*x*_), and substituting *τ*^*\**^ with the full goal Eq. (14) into the linearized body dynamics gives

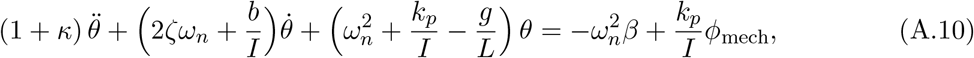

a second-order linear system whose equilibrium is Eq. (A.8). The acceleration-belief term raises the effective inertia by the factor 1+*κ* (1.8 at the baseline), and the passive damping adds *b/I* to the goal damping 2*ζω*_*n*_ (0.03 versus 8 s^−1^). The damping coefficient is positive, so the equilibrium is stable exactly when the closed-loop stiffness 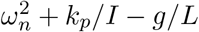is positive, that is, when 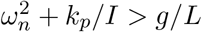. The denominator of *A* is this closed-loop stiffness, which is why the equilibrium expression and the stability requirement share the same term. This explains why a restorative generalized goal was required for standing at *k*_*p*_ = 0, and why intrinsic stiffness of 0.9 *mgL* restores stability at *ω*_*n*_ = 3.0 (Section 5.8). The slower root of Eq. (A.10) has time constant *τ*_slow_ of 1.0 s at the baseline, 2.6 s at *ω*_*n*_ = 3.5, and 0.45 s at *k*_*p*_ = 0.9 *mgL*; this value was used to set the perturbation hold (Appendix A.5). It is a linearized, fast-belief estimate; near the boundary both approximations degrade, and the condition *ω*_*n*_ = 3.2 (*τ*_slow_ = 14.7 s) was excluded from the main run because 10 *τ*_slow_ exceeded the hold cap.

### A.2 Reduction ratios

For a single perturbed channel *t* with perturbation amplitude *a*, Eq. (A.1) gives |*β*| = (*λ*_*t*_*/*Σ*λ*) *a*, and Eq. (A.9) maps this bias to the posture shift |*θ*_late_| = *A* (*λ*_*t*_*/*Σ*λ*) *a*. If the target-channel precision changes from *λ*_*t*_ to 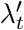(with precision sums Σ*λ* and Σ*λ*^*′*^) while the perturbation amplitude and the body and controller parameters are unchanged, the remaining response fraction is

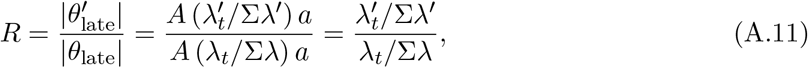

where 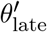 is the late posture under the changed precision: the common amplification factor *A* and the amplitude *a* cancel, so condition-wise reductions are determined by relative precision ratios alone. For the single-channel visual or proprioceptive manipulation,

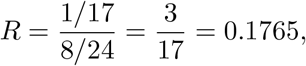

corresponding to an 82.35% reduction. In the equal-amplitude visual-proprioceptive conflict condition (*o* = +5°, *ϕ* = −5°), the two perturbation terms in Eq. (A.1) have the same sign, so the perturbed-channel weights add, giving

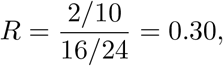

corresponding to a 70.00% reduction. These are linearized predictions; the simulated reductions reported in Section 5 are 82.3% and 69.8%, the small differences arising from the sin *θ* nonlinearity at the larger equilibrium angles. The sensitivity to the evaluation window is assessed separately in Appendix A.5.

For a physical support tilt, *θ*_late_ follows Eq. (A.8) with *β* = −*λ*_prop_*ϕ/*Σ*λ* and *ϕ*_mech_ = *ϕ*, so 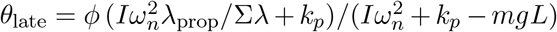 and the remaining fraction under a change of *λ*_prop_

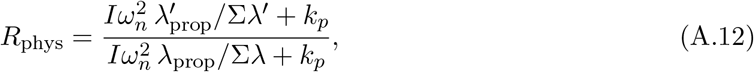

which equals Eq. (A.11) at *k*_*p*_ = 0 and tends to 1 as *k*_*p*_ *→ ∞*. For the 8-to-1 manipulation with *k*_*p*_ = 0.9 *mgL* = 618 N m rad^−1^, *R*_phys_ = (1120*/*17 + 618)*/*(1120*/*3 + 618) = 0.690.

### A.5 Numerical convergence, settling, and range of verification

Fig. A.1 compares every settled trial with the nonlinear equilibrium of Eq. (A.7). The maximum absolute residual over the 111 settled trials is 0.00056° (largest for *L* = 0.8 m, where |*θ*_late_| = 7.1°); residuals below 10^−9^ degrees did not occur. The residual from the linear prediction grows with |*θ*_late_| as expected from the sin *θ* correction and exceeds 0.01° only above about 4°. The numerical-convergence block (Fig. A.1C) shows that the late posture is unchanged, to within 2 *×*10^−5^ degrees, by the number of D-step iterations (16, 32, 64) and by halving the integration step (16 iterations at 2.5 ms and 32 at 5 ms apply the same number of belief updates per simulated second); this establishes that the steady values do not depend on the discretization, but not that every transient is discretization-independent. Fig. A.1D compares two evaluation windows: over 5 s to 6 s, the same trials deviate from the equilibrium by up to 0.3° (about 5% of the response), whereas the 14 s to 16 s window deviates by less than 0.001°. The slow closed-loop mode of Eq. (A.10) has a time constant of about 1 s at the baseline, so the short window contained a 3–4 time-constant residual.

**Figure A.1:**
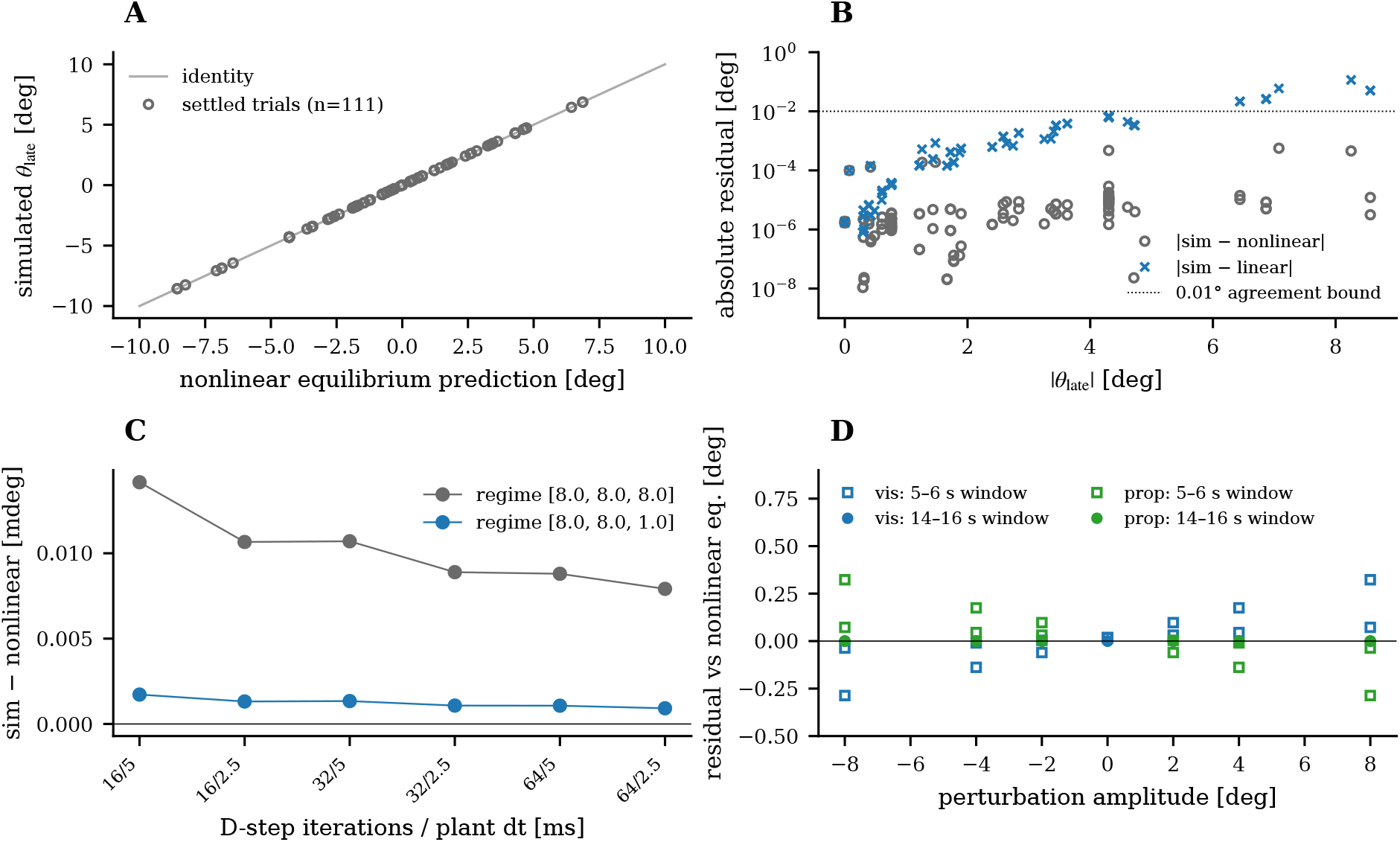
Steady-state verification. (A) Simulated late posture versus the nonlinear equilibrium for the 111 settled trials. (B) Absolute residual from the nonlinear equilibrium (circles) and from the linear prediction (crosses) versus |*θ*_late_|; dotted line, the 0.01° agreement bound. (C) Residual from the nonlinear equilibrium for the numerical-convergence block (D-step iterations / integration step in ms). (D) Residual of the 5 s to 6 s window (open squares) and of the 14 s to 16 s window (filled circles) for the perturbation-amplitude block.

The settling flags defined in Section 5 and the hold rule of Section 3.8 were specified before the main run: the hold was 14 s by default and 10 *τ*_slow_ for conditions with *τ*_slow_ *≥* 1.5 s (26 s at *ω*_*n*_ = 3.5, 19 s at *L* = 0.8 m), with a cap of 60 s. Fig. A.2 lists every trial of the main run with its flags; the condition *ω*_*n*_ = 3.2 (*τ*_slow_ = 14.7 s) was excluded by the cap and is shown as excluded. The 115 trials are discrete verification points chosen to vary one factor at a time; they do not map the stable region, which is given analytically by Eq. (A.10).

### A.6 Time course around a precision switch

Fig. A.3 expands 200 ms around the 8-to-1 switch of Fig. 7, with every 5 ms sample marked. The target channel’s gradient changes at the switch step; the tilt belief then changes over several numerical update steps while the body angle changes slowly; and the goal acceleration and torque follow the belief. The speed of the belief change depends on the numerical D-step setting (32 iterations per 5 ms step) and is not a physiological processing time. Table A.2 lists the 10–90% response times of body tilt for each transition, defined as follows: the start value is the posture sample at the event time, the end value is the late-window posture of the control trial with the corresponding constant precision (0 for the perturbation release), and *t*_10_ and *t*_90_ are the first samples after the event at which the normalized excursion reaches 0.1 and 0.9, without interpolation.

**Figure A.2:**
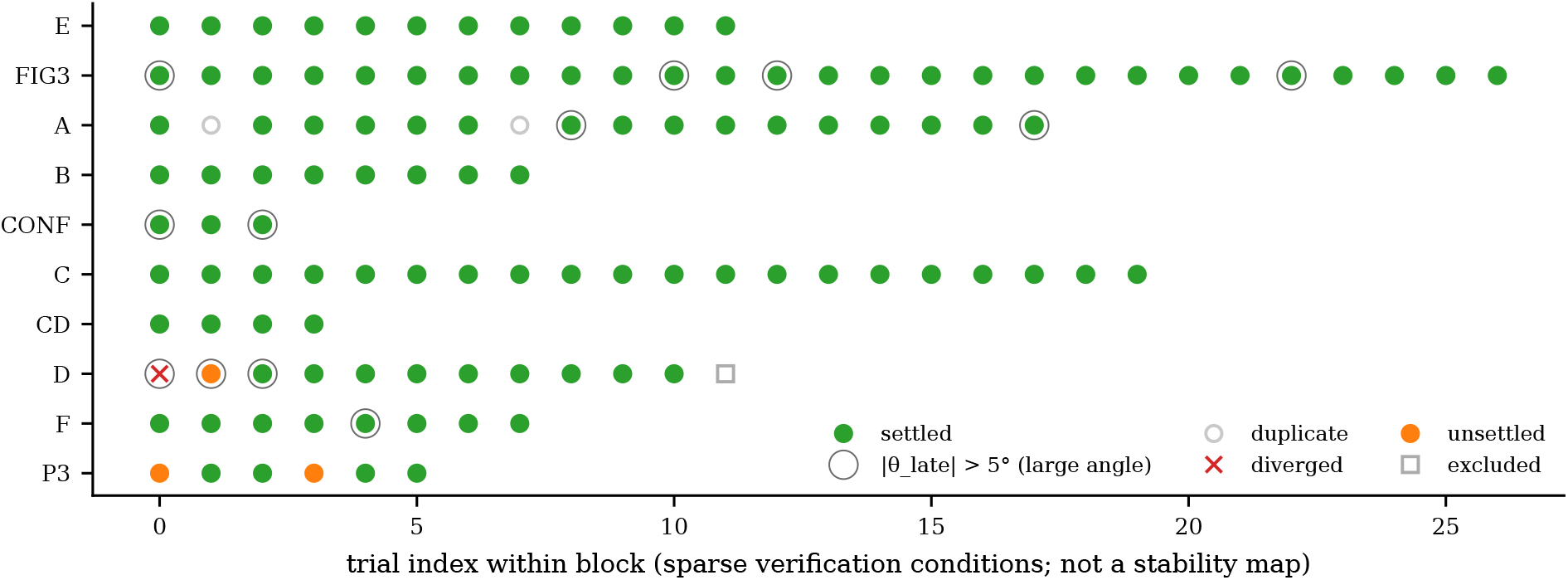
Trial-level flags of the main run, by block (E, numerical convergence; FIG3, perturbation amplitude; A, target precision; B, global scaling and global-low; CONF, combined conflict; C, intrinsic stiffness; CD, passive damping; D, goal dynamics; F, body scale; P3, precision switch). Filled green, settled; orange, finite and not diverged but not settled (none of these saturated); red cross, diverged; open square, excluded by the hold cap; open grey, duplicate specification reusing a result; large ring, |*θ*_late_| *>* 5°.

**Table A.2:** 10–90% response times of body tilt in the precision-switch trials (model indices under an imposed precision schedule; not physiological time constants).

| Channel | Event | Start [deg] | End [deg] | *t*<sub>10</sub> [s] | *t*<sub>90</sub> [s] |
|---|---|---|---|---|---|
| Visual | 8 → 1 at 7 s | −4.26 | −0.76 | 0.295 | 2.650 |
| Visual | 1 → 8 at 12 s | −0.79 | −4.30 | 0.290 | 2.655 |
| Visual | release at 17 s | −4.27 | 0 | 0.290 | 2.665 |
| Proprioceptive | 8 → 1 at 7 s | +4.26 | +0.76 | 0.295 | 2.650 |
| Proprioceptive | 1 → 8 at 12 s | +0.79 | +4.30 | 0.290 | 2.655 |
| Proprioceptive | release at 17 s | +4.27 | 0 | 0.290 | 2.665 |

The start samples are not fully settled states, because each segment lasts 5 s; no onset-latency index is defined.

### A.7 Robustness to stochastic observation noise

The main simulations used deterministic observations so that the effect of precision weighting could be isolated. To test whether the same reweighting pattern persists when precision is paired with actual observation-noise variance, we ran an additional stochastic robustness check. Zero-mean Gaussian noise was added to each generalized observation component with

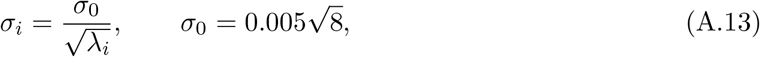

so that 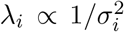. This check was intended as a robustness test of the precision–variance correspondence, not as a fitted physiological sensor-noise model. We used the representative +5° visual-offset and sensory-only support-tilt perturbations with *k*_*p*_ = 0, 20 random seeds per condition, and the same long-hold protocol and late response window as in the main analyses. The deterministic reference for each condition is the main-run trial under the identical protocol. For the visual and proprioceptive target channels the order-0 gradient has an angle component only, so the contribution index *C*_*i*_ of Eq. (25) equals the Euclidean norm of the order-0 gradient for those channels; this identity is not claimed for the vestibular channel.

**Figure A.3:**
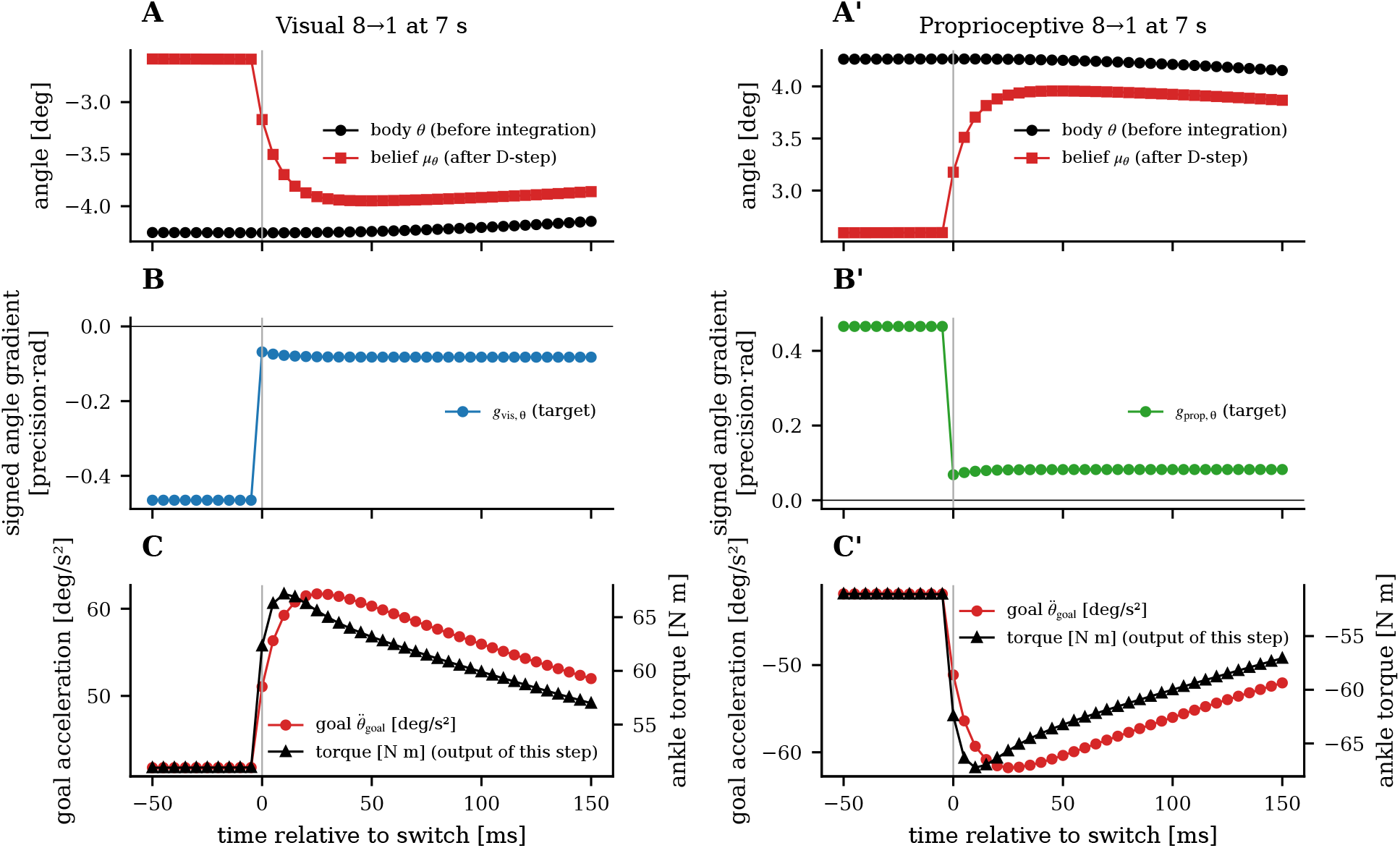
200 ms around the 8-to-1 switch at 7 s for the visual (left) and proprioceptive (right) switch trials; markers are the 5 ms samples. (A) Body tilt (before integration) and tilt belief (after the D-step). (B) Signed angle gradient of the target channel. (C) Preferred acceleration and ankle torque (output of the step).

All 100 stochastic runs satisfied the stability criteria (Table A.3); the deterministic settling flags were recorded but not applied, because the angular velocity does not vanish under noise. Computed from the condition means, visual precision reduction decreased the visual-offset posture shift by 82.29% and the visual state-update contribution by 82.35%, and proprioceptive precision reduction decreased the support-tilt posture shift by 82.31% and the proprioceptive contribution by 82.38% (deterministic: 82.33%). The global low-precision condition reduced the target-channel contribution by 87.5% while changing the mean posture shift by −0.08% relative to equal precision; that is, it did not show the bias reduction that the selective manipulation showed. Thus, pairing the precision parameters with matched observation-noise variances preserved the deterministic reweighting pattern under the same protocol.

**Table A.3:** Stochastic observation-noise robustness check under the long-hold protocol. Values are mean ± sample SD across 20 seeds; the deterministic value is the main-run trial with the same parameters. Seed-to-seed variation reflects the imposed noise, not individual differences.

| Condition | Stable runs | θ<sub>late</sub> [deg] | Deterministic [deg] | Target-channel *C<sub>i</sub>* |
|---|---|---|---|---|
| Visual equal (8, 8, 8) | 20/20 | −4.304 ± 0.021 | −4.302 | 0.4654 ± 0.0016 |
| Visual low (8, 8, 1) | 20/20 | −0.762 ± 0.024 | −0.760 | 0.0821 ± 0.0007 |
| Proprioceptive equal (8, 8, 8) | 20/20 | +4.299 ± 0.021 | +4.302 | 0.4647 ± 0.0018 |
| Proprioceptive low (8, 1, 8) | 20/20 | +0.760 ± 0.022 | +0.760 | 0.0819 ± 0.0007 |
| Visual global low (1, 1, 1) | 20/20 | −4.308 ± 0.057 | −4.302 | 0.0582 ± 0.0006 |

## References

Adams, R. A., Shipp, S., and Friston, K. J. (2013). Predictions not commands: active inference in the motor system. Brain Structure and Function, 218(3):611–643.

Assländer, L. and Peterka, R. J. (2014). Sensory reweighting dynamics in human postural control. Journal of Neurophysiology, 111(9):1852–1864.

Assländer, L. and Peterka, R. J. (2016). Sensory reweighting dynamics following removal and addition of visual and proprioceptive cues. Journal of Neurophysiology, 116(2):272–285.

Baltieri, M. and Buckley, C. L. (2019). PID control as a process of active inference with linear generative models. Entropy, 21(3):257.

Buckley, C. L., Kim, C. S., McGregor, S., and Seth, A. K. (2017). The free energy principle for action and perception: A mathematical review. Journal of Mathematical Psychology, 81:55–79.

Carver, S., Kiemel, T., and Jeka, J. J. (2006). Modeling the dynamics of sensory reweighting. Biological Cybernetics, 95(2):123–134.

Chiba, R., Takakusaki, K., Ota, J., Yozu, A., and Haga, N. (2016). Human upright posture control models based on multisensory inputs; in fast and slow dynamics. Neuroscience Research, 104:96–104.

Ciria, A., Schillaci, G., Pezzulo, G., Hafner, V. V., and Lara, B. (2021). Predictive processing in cognitive robotics: a review. Neural Computation, 33(5):1402–1432.

Ernst, M. O. and Banks, M. S. (2002). Humans integrate visual and haptic information in a statistically optimal fashion. Nature, 415(6870):429–433.

Feldman, H. and Friston, K. J. (2010). Attention, uncertainty, and free-energy. Frontiers in Human Neuroscience, 4:215.

Friston, K. (2010). The free-energy principle: a unified brain theory? Nature Reviews Neuroscience, 11(2):127–138.

Friston, K., Daunizeau, J., Kilner, J., and Kiebel, S. J. (2010). Action and behavior: a free-energy formulation. Biological Cybernetics, 102(3):227–260.

Friston, K. J., Trujillo-Barreto, N., and Daunizeau, J. (2008). DEM: A variational treatment of dynamic systems. NeuroImage, 41(3):849–885.

Honeine, J.-L., Crisafulli, O., Sozzi, S., and Schieppati, M. (2015). Processing time of addition or withdrawal of single or combined balance-stabilizing haptic and visual information. Journal of Neurophysiology, 114(6):3097–3110.

Hwang, S., Agada, P., Kiemel, T., and Jeka, J. J. (2016). Identification of the unstable human postural control system. Frontiers in Systems Neuroscience, 10:22.

Jeka, J., Kiemel, T., Creath, R., Horak, F., and Peterka, R. (2004). Controlling human upright posture: velocity information is more accurate than position or acceleration. Journal of Neurophysiology, 92(4):2368–2379.

Kanai, R., Komura, Y., Shipp, S., and Friston, K. (2015). Cerebral hierarchies: predictive processing, precision and the pulvinar. Philosophical Transactions of the Royal Society B, 370(1668):20140169.

Kiemel, T., Oie, K. S., and Jeka, J. J. (2002). Multisensory fusion and the stochastic structure of postural sway. Biological Cybernetics, 87:262–277.

Knill, D. C. and Pouget, A. (2004). The Bayesian brain: the role of uncertainty in neural coding and computation. Trends in Neurosciences, 27(12):712–719.

Körding, K. P. and Wolpert, D. M. (2004). Bayesian integration in sensorimotor learning. Nature, 427:244–247.

Lanillos, P., Meo, C., Pezzato, C., Meera, A. A., Baioumy, M., Ohata, W., Tschantz, A., Millidge, B., Wisse, M., Buckley, C. L., and Tani, J. (2021). Active inference in robotics and artificial agents: survey and challenges. arXiv.

Le Mouël, C. and Brette, R. (2019). Anticipatory coadaptation of ankle stiffness and sensorimotor gain for standing balance. PLOS Computational Biology, 15(11):e1007463.

Limanowski, J. (2022). Precision control for a flexible body representation. Neuroscience & Biobehavioral Reviews, 134:104401.

Limanowski, J. and Friston, K. (2020). Active inference under visuo-proprioceptive conflict: Simulation and empirical results. Scientific Reports, 10(1):4010.

Lippi, V. and Mergner, T. (2017). Human-derived disturbance estimation and compensation (DEC) method lends itself to a modular sensorimotor control in a humanoid robot. Frontiers in Neurorobotics, 11:49.

Loram, I. D., Gollee, H., van de Kamp, C., and Gawthrop, P. J. (2022). Is intermittent control the source of the non-linear oscillatory component (0.2–2 Hz) in human balance control? IEEE Transactions on Biomedical Engineering, 69(12):3623–3634.

Loram, I. D. and Lakie, M. (2002). Direct measurement of human ankle stiffness during quiet standing: the intrinsic mechanical stiffness is insufficient for stability. The Journal of Physiology, 545(3):1041–1053.

Mergner, T. (2010). A neurological view on reactive human stance control. Annual Reviews in Control, 34(2):177–198.

Morasso, P., Cherif, A., and Zenzeri, J. (2019). Quiet standing: The single inverted pendulum model is not so bad after all. PLOS ONE, 14(3):e0213870.

Nashner, L. M., Black, F. O., and Wall, C. (1982). Adaptation to altered support and visual conditions during stance: patients with vestibular deficits. The Journal of Neuroscience, 2(5):536–544.

Oie, K. S., Kiemel, T., and Jeka, J. J. (2002). Multisensory fusion: simultaneous re-weighting of vision and touch for the control of human posture. Cognitive Brain Research, 14(1):164–176.

Oliver, G., Lanillos, P., and Cheng, G. (2022). An empirical study of active inference on a humanoid robot. IEEE Transactions on Cognitive and Developmental Systems, 14(2):462–471.

Parr, T., Limanowski, J., Rawji, V., and Friston, K. (2021). The computational neurology of movement under active inference. Brain, 144(6):1799–1818.

Parr, T., Pezzulo, G., and Friston, K. J. (2022). Active Inference: The Free Energy Principle in Mind, Brain, and Behavior. MIT Press.

Pasma, J. H., Engelhart, D., Maier, A. B., Schouten, A. C., van der Kooij, H., and Meskers, C. G. M. (2015). Changes in sensory reweighting of proprioceptive information during standing balance with age and disease. Journal of Neurophysiology, 114(6):3220–3233.

Peterka, R. J. (2002). Sensorimotor integration in human postural control. Journal of Neurophysiology, 88(3):1097–1118.

Peterka, R. J. and Loughlin, P. J. (2004). Dynamic regulation of sensorimotor integration in human postural control. Journal of Neurophysiology, 91(1):410–423.

Peterka, R. J., Murchison, C. F., Parrington, L., Fino, P. C., and King, L. A. (2018). Implementation of a central sensorimotor integration test for characterization of human balance control during stance. Frontiers in Neurology, 9:1045.

Pio-Lopez, L., Nizard, A., Friston, K., and Pezzulo, G. (2016). Active inference and robot control: a case study. Journal of The Royal Society Interface, 13(122):20160616.

Priorelli, M., Maggiore, F., Maselli, A., Donnarumma, F., Maisto, D., Mannella, F., Stoianov, I. P., and Pezzulo, G. (2024). Modeling motor control in continuous time active inference: a survey. IEEE Transactions on Cognitive and Developmental Systems, 16(2):485–500.

Rasman, B. G., Forbes, P. A., Peters, R. M., Ortiz, O., Franks, I., Inglis, J. T., Chua, R., and Blouin, J.-S. (2021). Learning to stand with unexpected sensorimotor delays. eLife, 10:e65085.

Sakanaka, T. E., Gill, J., Lakie, M. D., and Reynolds, R. F. (2018). Intrinsic ankle stiffness during standing increases with ankle torque and passive stretch of the Achilles tendon. PLOS ONE, 13(3):e0193850.

Sakanaka, T. E., Lakie, M., and Reynolds, R. F. (2021). Individual differences in intrinsic ankle stiffness and their relationship to body sway and ankle torque. PLOS ONE, 16(1):e0244993.

Stevenson, I. H., Fernandes, H. L., Vilares, I., Wei, K., and Körding, K. P. (2009). Bayesian integration and non-linear feedback control in a full-body motor task. PLoS Computational Biology, 5(12):e1000629.

Tisserand, R., Dakin, C. J., Van der Loos, M. H. F., Croft, E. A., Inglis, T. J., and Blouin, J.-S. (2018). Down regulation of vestibular balance stabilizing mechanisms to enable transition between motor states. eLife, 7:e36123.

Todorov, E. and Jordan, M. I. (2002). Optimal feedback control as a theory of motor coordination. Nature Neuroscience, 5(11):1226–1235.

van der Kooij, H., Jacobs, R., Koopman, B., and van der Helm, F. (2001). An adaptive model of sensory integration in a dynamic environment applied to human stance control. Biological Cybernetics, 84:103–115.

van Kordelaar, J., Pasma, J. H., Cenciarini, M., Schouten, A. C., van der Kooij, H., and Maurer, C. (2018). The reliance on vestibular information during standing balance control decreases with severity of vestibular dysfunction. Frontiers in Neurology, 9:371.

Winter, D. A., Patla, A. E., Prince, F., Ishac, M., and Gielo-Perczak, K. (1998). Stiffness control of balance in quiet standing. Journal of Neurophysiology, 80(3):1211–1221.

